# Ipl1-controlled attachment maturation regulates Mps1 association with its kinetochore receptor

**DOI:** 10.1101/2023.10.30.564738

**Authors:** Richard Pleuger, Christian Cozma, Simone Hohoff, Christian Denkhaus, Alexander Dudziak, Farnusch Kaschani, Andrea Musacchio, Ingrid R. Vetter, Stefan Westermann

## Abstract

Faithful chromosome segregation requires that sister chromatids establish bi-oriented kinetochore-microtubule attachments. The spindle assembly checkpoint (SAC) prevents premature anaphase onset with incomplete attachments. However, how microtubule attachment and checkpoint signaling are coordinated remains unclear. The conserved kinase Mps1 initiates SAC signaling by localizing transiently to kinetochores in prometaphase and is released upon bi-orientation. Using biochemistry, structure predictions, and cellular assays, we shed light on this dynamic behaviour in *Saccharomyces cerevisiae*. A conserved N-terminal segment of Mps1 binds the neck region of Ndc80:Nuf2, the main microtubule receptor of kinetochores. Mutational disruption of this interface, located at the backside of the paired CH-domains and opposite the microtubule-binding site, prevents Mps1 localization, eliminates SAC signalling, and impairs growth. The same interface of Ndc80:Nuf2 binds the microtubule-associated Dam1 complex. We demonstrate that the error correction kinase Ipl1/Aurora B controls the competition between Dam1 and Mps1 for the same binding site. Thus, binding of the Dam1 complex to Ndc80:Nuf2 may release Mps1 from the kinetochore to allow anaphase onset.

## Introduction

Chromosome segregation during cell division in eukaryotes requires that sister chromatids are attached to opposite poles of the mitotic spindle, allowing their subsequent segregation to the two daughter cells. This configuration, termed bi-orientation, is gradually achieved during prometaphase of mitosis through the activity of kinetochores, large protein complexes that link chromosomes to spindle microtubules. Kinetochores also ensure that as long as bi-orientation has not been successfully achieved for all chromosomes, the onset of anaphase is prevented by inhibiting the E3 Ubiquitin-ligase Anaphase Promoting Complex/Cyclosome (APC/C) through a signaling cascade termed the spindle assembly checkpoint (SAC)^1,2^. Genetic screens in yeast identified the main components of the SAC, the *mad* and *bub* genes^3,4^. In addition to these core checkpoint genes, the conserved protein kinase Mps1 (monopolar spindles-1), plays a crucial role in the SAC and the establishment of bi-oriented attachments ^5–7^. Mps1 is at or near the top of the checkpoint signaling cascade. This is most clearly demonstrated by the observation that overexpression of Mps1 from a Galactose-inducible promoter arrests yeast cells permanently in metaphase with fully formed bi-oriented attachments. This metaphase arrest is dependent on the core checkpoints genes^8^. During checkpoint signaling, Mps1 contributes to the initial recruitment of Bub1:Bub3 through phosphorylation of the kinetochore subunit Spc105/Knl1^9–11^. It also contributes directly to the catalytic generation of a diffusible inhibitor of the APC/C, the mitotic checkpoint complex (MCC)^12–14^. Previous biochemical and genetic experiments indicated that Mps1 is linked to the outer kinetochore KMN network, most likely through a direct interaction with the main microtubule receptor of the kinetochore, the Ndc80 complex (Ndc80c)^15^. Apart from its role in the SAC, Mps1 also contributes to bi-orientation, at least in part by phosphorylating the N-terminal tail of Ndc80^16^. Finally, Mps1 is also essential for spindle pole body duplication^17^ in budding yeast, complicating the analysis of its specific contributions to kinetochore function.

After successful bi-orientation, Mps1 phosphorylation is suppressed and substrates are dephosphorylated to silence the SAC^18,19^. It is expected that control of Mps1 recruitment to, and release from, kinetochores, may be crucial for this dynamic regulation. Thus, how Mps1 is recruited to kinetochores and the regulation of its activity there constitute key questions in the field. Recent studies put forward different models to explain how Mps1 kinetochore occupancy is related to the attachment configuration and achievement of bi-orientation. In the “competition model”, microtubule-binding by Ndc80 is mutually exclusive with the recruitment of Mps1. This model is based on in-vitro biochemical experiments demonstrating that binding of Mps1 to the CH-domains of human Ndc80 is competitive with microtubule binding^20,21^. A call for further scrutiny of the competition model, however, comes from observations that Mps1 can be induced to become re-recruited to kinetochores even when Ndc80 remains bound to microtubules^22,23^. In the “spatial separation model”, on the other hand, establishment of tension during bi-orientation separates Mps1 from key substrates such as Spc105/Knl1, possibly involving a barrier function of other kinetochore components. This model is inspired by artificial recruitment experiments in budding yeast and it remains unclear whether it fully reflects the natural mode of Mps1 regulation^24^.

A fundamental limitation in our understanding of Mps1 regulation at the kinetochore is that the binding interface between Mps1 and Ndc80c has not been fully defined in any system. For instance, Mps1 autophosphorylation has also been suggested as a main driving force for regulation of recruitment^25^, as it limits the amount of Mps1 at kinetochores. At present, however, it remains unclear whether this mechanism contributes to Mps1 regulation during an authentic cell cycle, in part because neither the binding interface, nor the relevant phosphorylation sites have been fully described. To shed light on this regulation, it is therefore essential that the kinetochore binding sites for Mps1 are identified. Here we establish the binding mechanism between Mps1 and Ndc80c and present separation-of-function mutants that allow us to probe different mechanisms of Mps1 recruitment regulation. The finding that the Ndc80c:Mps1 interface coincides with the binding site of Dam1 C-terminus illuminates the mechanistic relationship between the error correction machinery and the spindle assembly checkpoint. We show that Ipl1-controlled error correction, mediated through phosphorylation of Dam1, promotes Mps1 binding by avoiding competition between Mps1 and Dam1 for Ndc80c, allowing sustained checkpoint signaling in prometaphase. The maturation of kinetochore end-on attachments is then coupled to SAC silencing via Dam1-dependent displacement of Mps1 from its kinetochore receptor binding site.

## Results

### An Mps1-Ndc80c binding interface at the CH domain neck

Previous genetic experiments in budding yeast identified the N-terminal part of Mps1 as being important for chromosome segregation^26^. To determine if Mps1 interacts directly with Ndc80c, and if so, which segment of Mps1 is required, we fused N-terminal Mps1 segments to Maltose-binding protein (MBP) and performed pull-down assays with Baculovirus-expressed Ndc80 complex carrying a Flag-tag at the Carboxy-terminus of the Ndc80 subunit. We found that a fusion encompassing Mps1 residues 1–173 efficiently pulled down Ndc80c-Flag from Sf9 cell lysates, while Mps1 1–143 or shorter N-terminal segments failed to do so (Figure 1 B). Thus, Mps1 segment 143-173 contributes critically to Ndc80c binding, likely explaining the meiotic chromosome segregation defects of an Mps1^R170S^ mutant^27^.

**Figure 1:**
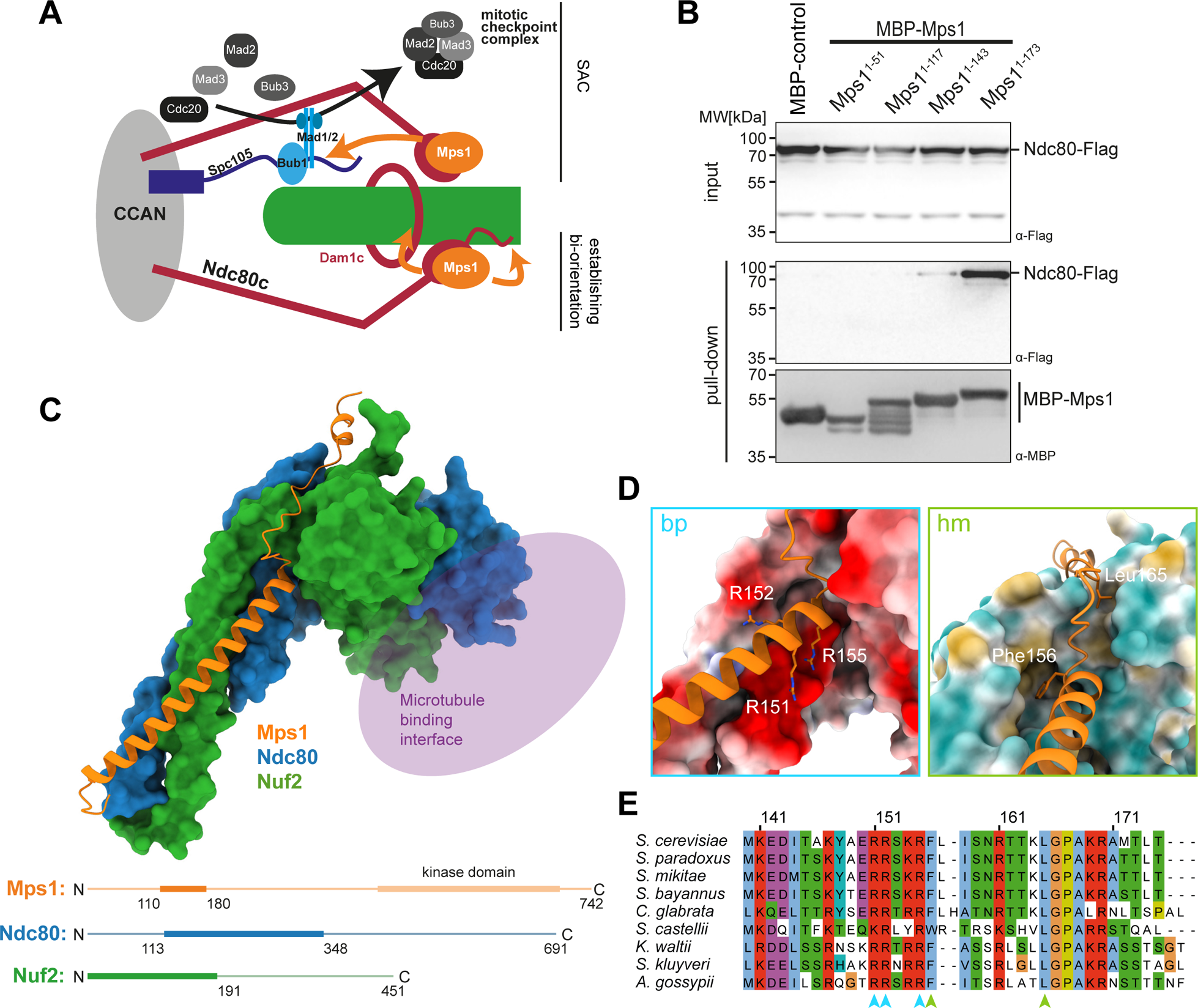
Mps1 engages with the Ndc80c on the backside of the CH-domains. A: Schematic illustration illustrating Mps1 functions at the kinetochore. Ndc80 acts as the main kinetochore receptor for Mps1. Mps1 promotes mitotic checkpoint complex assembly as well as establishment of bi-oriented end-on attachments. B: Pull-down assay using MBP fused to Mps1 N-terminal fragments as bait and lysates from SF9 cells infected to express Flag tagged Ndc80 complex. Only MBP-Mps1^1–180^ binds Ndc80c efficiently. C: Alphafold2 structure prediction of a trimeric complex Mps1^110–180^ (orange) :Ndc80^113–348^ (blue) :Nuf2^1–191^ (green). Mps1 forms a three helix bundle with the coiled-coils of Ndc80 and Nuf2. An unstructured segment winds around the neck of the CH-domains opposite of the microtubule binding interface. D: Interaction surfaces between Mps1 and the Ndc80:Nuf2 dimer predicted by a PISA analysis. Highlighted residues in Mps1 were targeted for point mutations in this study. Blue: basic patch mutant “bp”; Green: hydrophobic mutant “hm”. E: Multiple sequence alignment of Mps1 proteins in different yeasts spanning the main interacting interface. Arrows below indicate the residues mutated for the “bp” (blue) or the “hm” (green) mutant.

Based on this preliminary delineation of the binding motif we performed AF2 structure predictions with the N-terminal Mps1 segment and the N-terminal, two-subunit Ndc80:Nuf2 complex as inputs. AF2-Multimer (version 2.3.1)^28,29^ models predicted with high confidence an interaction mode in which residues 111-170 of Mps1 engage with Ndc80:Nuf2 as a helical bundle at the “neck” region of Ndc80:Nuf2, just underneath the paired CH-domains (Figure 1C, Supporting Figure 1). Following a predicted alpha-helical segment (residues 111-156), the disordered Mps1 residues 157–170 drape across the surface of the tandem CH-domains, opposite to the microtubule-binding interface. A crystal structure of an Mps1 peptide bound to Ndc80c provides an experimental confirmation of the overall accuracy of the AF2 model (Zahm and Harrison, personal communication).

A PISA (Proteins, Interfaces, Structures and Assemblies) analysis^30^ on the AF2 model calculated that the interface between Mps1 and Nuf2 is larger than that between Mps1 and Ndc80 (1266 Å^2^ vs 680 Å^2^). The analysis also allowed to identify residues predicted to be crucial for binding. At the C-terminal end of the Mps1 helix, residues R151, R152 and R155 constitute a “basic patch” predicted to form ionic interactions with Ndc80:D265, Ndc80:E269, and Ndc80:E272 (interacting with Mps1:R151 and Mps1:R155) and Nuf2:E148 (interacting with Mps1:R152) (Figure 1D). Another crucial element of the interface is a network of hydrophobic interactions formed around Mps1:F156, which engages Ndc80:L279 and Nuf2:I144, and around Mps1:L165 (which engages Nuf2:S22, Nuf2:C23, and Nuf2:F52).

### The Mps1^hm^ mutant is defective in Ndc80c binding in vitro

Based on this analysis and the conservation of residues among different yeast species, we designed two sets of interface mutants (Figure 1D): a basic patch mutant comprising Mps1 R151D R152D R155D (bp) and a hydrophobic mutant (hm) comprising Mps1 F156D L165A. We also combined the mutations in a single mutant (hm+bp). We tested these mutants in the context of an Mps1 1–250 MBP fusion construct in pull-down assays with Ndc80c-Flag. Consistent with a critical function of Mps1:F156 and Mps1:L165 in the binding interface, the Mps1^hm^ and Mps1^hm+bp^ mutants displayed strongly reduced binding of Ndc80c-Flag from the Sf9 lysate. The Mps1^bp^ mutant, on the other hand, bound Ndc80c similarly to the wild-type Mps1, if not better (Figure 2A).

**Figure 2:**
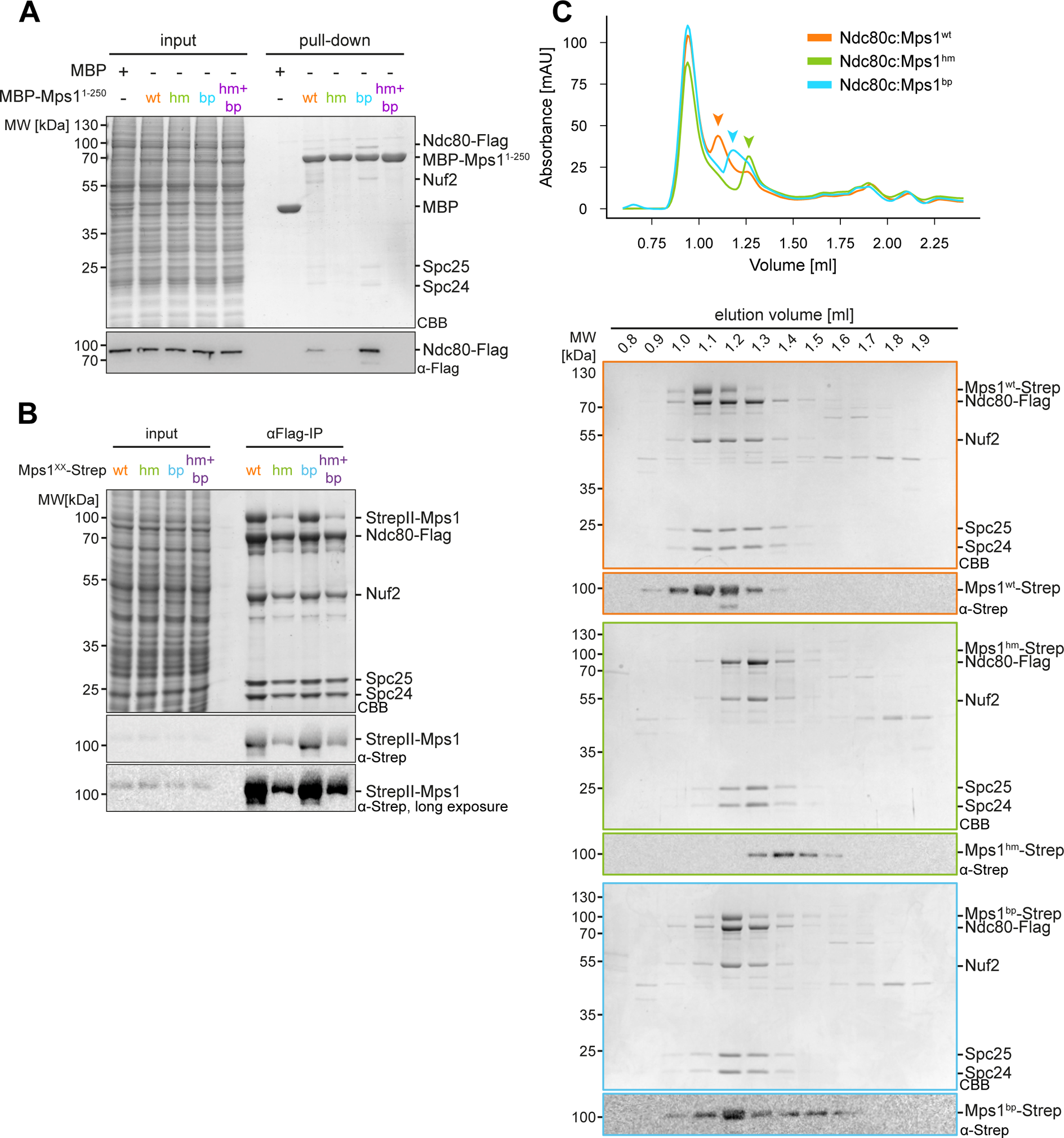
Interface point mutations destabilize the interaction between Mps1 and the Ndc80 complex *in vitro*. A: Pull-down assay showing the ability of MBP-Mps1^1–250^ (wt or interface mutants) to bind the Ndc80c. MBP-Mps1^1–250, hm^ and Mps1^1–250, hm+bp^ are defective in binding Ndc80c-Flag to the amylose resin. B: Co-immunoprecipitation of the Ndc80c (Ndc80-Flag) and Mps1-StrepII (wt or interface mutants) from Sf9 lysates after co-infection in SF9 cells. Mps1^wt^ and Mps1^bp^ are purified as an almost stochiometric complex. Mps1^hm^ and Mps1^hm+bp^ show drastically reduced levels in the IP elution. I: input, U: unbound, E: elution. C: Size exclusion chromatography runs of eluate samples of Co-immunoprecipitated StrepII-Mps1:Ndc80c from B. Arrow heads show the peaks corresponding to Ndc80c or StrepII-Mps1^XX^:Ndc80c if a the proteins run as a complex. The high peak at approximately 0.8 ml elution volume most likely results from DNA contaminations in the input samples. Column: Superose6 3.2/300.

We next analyzed the interaction with full-length proteins by co-infecting Sf9 cells with baculoviruses encoding Ndc80c-Flag and StrepII-tagged full-length Mps1 in wild-type or mutant form. SDS-PAGE and western blotting showed that wild-type and Mps1^bp^ co-purified in near-stoichiometric amounts with Ndc80c. By contrast, only little Mps1^hm^ or Mps1^hm+bp^ was detected in the elution fractions (Figure 2 B).

We further analyzed the isolated Ndc80c:Mps1 complexes by analytical size exclusion chromatography (SEC). Wild-type Mps1 formed a stable complex with Ndc80c, which eluted early from the column and was well separated from excess free Ndc80c (Figure 2C). The residual amount of Mps1^hm^ mutant which co-purified with Ndc80c, by contrast, was separated from the complex during SEC, indicative of impaired binding. The Mps1^bp^ mutant co-eluted with Ndc80c, but the complex peak was shifted relative to Ndc80c:Mps1^wt^, indicative of reduced binding affinity of the bp mutant for the Ndc80 complex (Figure 2C). Thus, our mutants validate the predicted binding interface *in vitro* and display different residual binding affinities for Ndc80c with Mps1^hm^ showing the most penetrant effects.

### Mps1 interface mutants impair kinetochore localization and viability in cells

To test the effects of Mps1-Ndc80c interface mutations in cells, we constructed yeast strains expressing Mps1-mNeonGreen (Mps1-mNG) in wild-type or mutant form from its endogenous promoter. A Nuf2-mCherry marker in the same strain allowed us to evaluate kinetochore enrichment by live cell microscopy (Figure 3A). We quantified Mps1 fluorescence intensity at kinetochores and plotted it as a function of cell cycle progression relative to anaphase onset. Wild-type Mps1-mNG displayed a dynamic kinetochore association, showing pronounced enrichment in pro-metaphase and then a steep decline just prior to anaphase onset (Figure 3B). Mps1 kinetochore levels then started to rise again about 50 min after anaphase onset as cells entered the next G1 phase. Quantification revealed that the Mps1 interface mutants fully prevented the kinetochore enrichment in prometaphase, despite being expressed at identical levels to the wild-type (Figure 3A, B, Supporting Figure 3A). This effect was specific to the kinetochore enrichment of Mps1 (calculated as the difference between the fluorescence intensities at kinetochores and in the nucleus), since a comparison of the nuclear background Mps1 levels between wild-type and mutants revealed no significant differences and showed a drop in levels from metaphase to anaphase for all tested Mps1 versions (Supporting Figure 3B). This result also shows that the reduction of Mps1 levels between metaphase to anaphase is independent of kinetochore association. Importantly, the intensities of Nuf2-mCherry were essentially identical in strains expressing wild-type or mutant Mps1, indicating that lack of kinetochore enrichment of the Mps1 mutants is not caused by reduced levels of the receptor, Ndc80c (Supporting Figure 3C). We conclude that the generated interface mutants prevent Mps1 kinetochore enrichment in cells.

**Figure 3:**
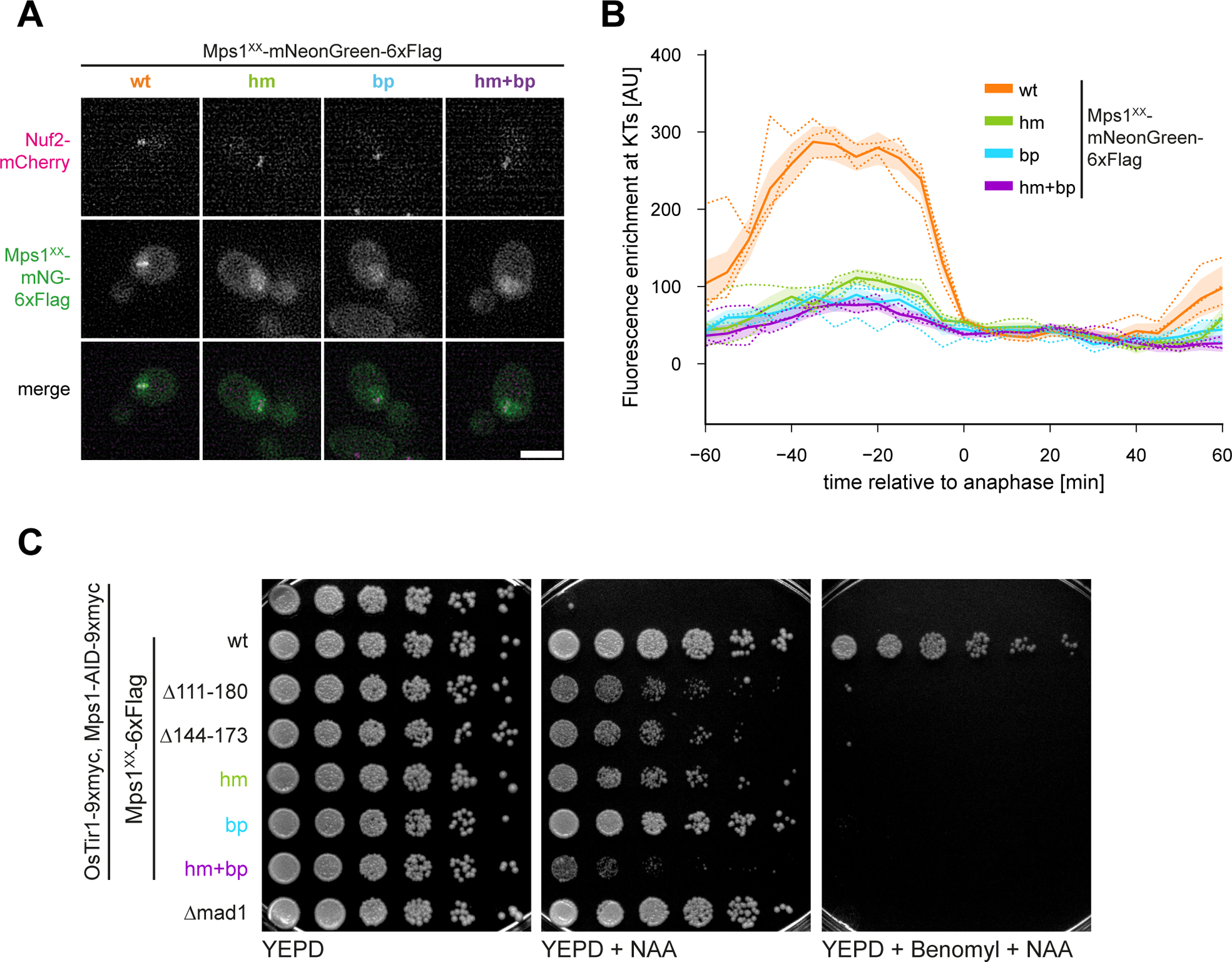
Mps1 interface mutants cannot localize to the kinetochore *in vivo*. A: Representative micrographs of metaphase cells expressing Nuf2-mCherry and Mps1-mNeonGreen-6xFlag (wt or interface mutants). Mps1^wt^-mNeonGreen shows a focused signal co-localized with Nuf2-mCherry signals on short spindles. The Mps1 interface mutants show a broader circular signal corresponding to the nuclear background. Scale bar: 5 μm B: Quantification of the kinetochore enrichment (kinetochore fluorescence intensity minus nuclear background intensity) of Mps1-mNeonGreen-6xFlag (wt and interface mutants). Values are plotted relative to anaphase onset, defined as the continuous separation of the kinetochore clusters. Mean and SEM of all replicates are indicated as a continuous line with borders, the means of three replicates are indicated as dotted lines. C: Serial dilution assay indicating growth phenotype of conditional Mps1 alleles on YEPD, YEPD + 1 mM NAA or YEPD + 1 mM NAA + 20 μg/ml benomyl. Except for Mps1^bp^, mutations or deletions in the segment of Mps1 that binds the Ndc80 complex display growth defects of different severity on rich media supplemented with NAA. On plates supplemented with NAA and benomyl, only the wildtype allele allows viability.

Mps1 has an essential function in spindle pole duplication^17^. It is therefore important to assess if the interface mutants presented here constitute true separation-of-function alleles. To test this, we fused an auxin-inducible degron (AID) tag to the endogenous Mps1 to acutely deplete the protein. In these cells we expressed Flag-tagged wild-type and mutant Mps1 rescue constructs from a marker locus. We optimized the AID system by placing OsTIR under a strong constitutive promoter, which allowed efficient depletion of endogenous Mps1 by the addition of NAA (Supporting Figure 3D). Serial dilution assays showed that the Mps1-AID strain was inviable in the presence of NAA, depending on both the AID tag and the expression of OsTIR, and could be rescued by expression of an Mps1 wild-type construct (Figure 3C and Supporting Figure 3E).

We used the Mps1-AID strain to assess if the Ndc80c-binding mutants we have identified support the essential requirement of Mps1 in spindle pole body duplication (SPB). For this, we labelled the SPB component Spc42 and analyzed SPB duplication in large-budded yeast cells 90 min after NAA treatment (Supporting Figure 4A). In the absence of a rescue allele, only 10% of NAA-treated large budded Mps1-AID cells contained two spindle pole bodies (Supporting Figure 4B and C). This defect was fully rescued not only by expression of Mps1^wt^, but also of Mps1^hm^, Mps1^bp^ or Mps1^hm+bp^ mutants. Thus, we have identified separation of function mutants of Mps1 that are fully proficient in bipolar spindle assembly and can be used to specifically interrogate the kinetochore function of Mps1.

**Figure 4:**
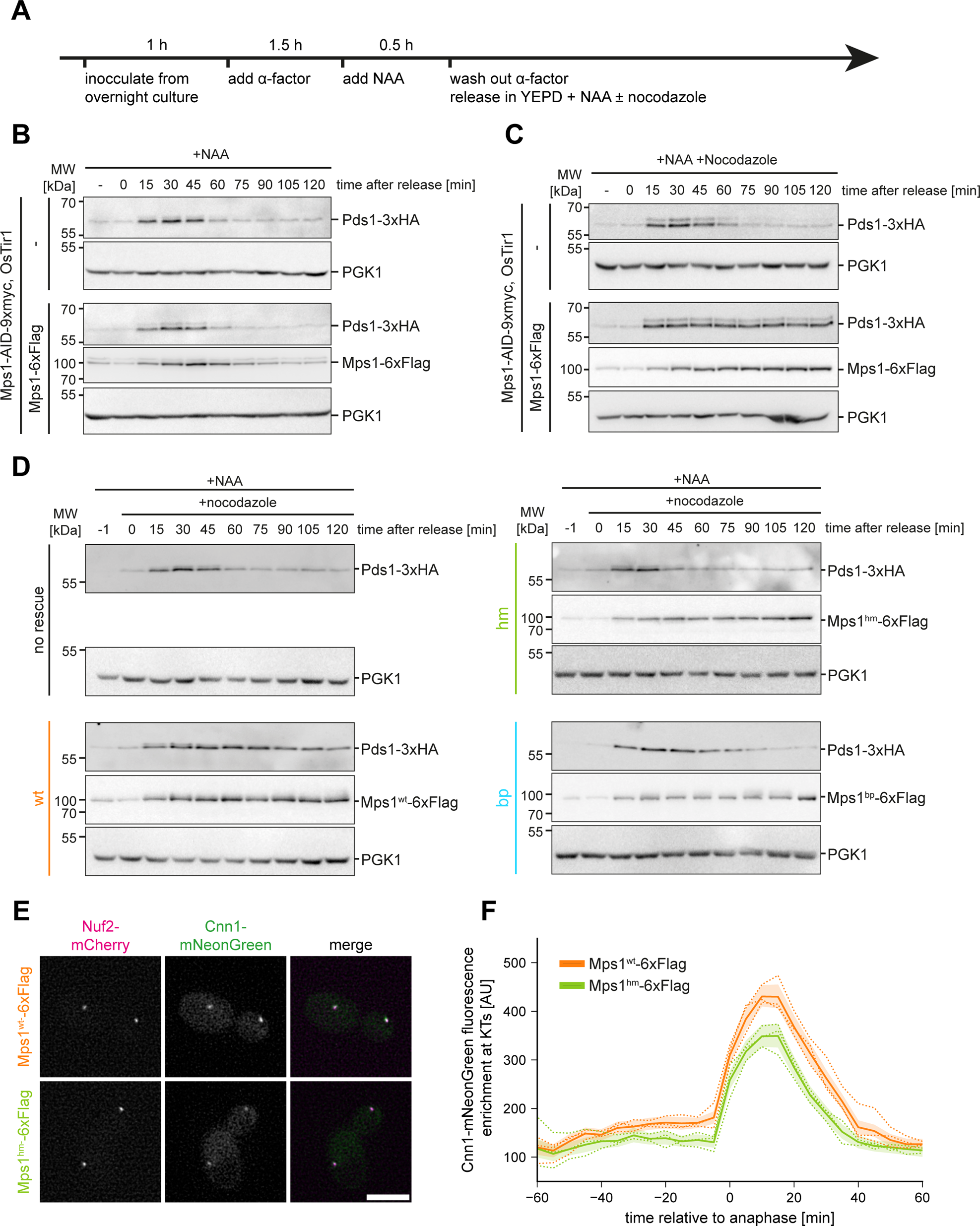
Interface mutants are defective in the kinetochore functions of Mps1. A: Schematic of the experimental design to test mitotic progression with or without nocodazole treatment in dependence of the tested Mps1 alleles. B: Cell cycle experiment of cells depleted of the endogenous Mps1 with or without an Mps1-6xFlag rescue allele. The presence of Mps1 is not determining the progression dynamics through the cell cycle in an otherwise unperturbed mitosis. C: Nocodazole arrest experiment of cells depleted of the endogenous Mps1 with or without an Mps1-6xFlag rescue allele. Cells depleted from the endogenous Mps1 cannot arrest in mitosis upon nocodazole treatment. Introduction of an exogenous Mps1 allele allows cells to stabilize Pds1 upon nocodazole treatment. D: Nocodazole arrest experiment of cells depleted of the endogenous Mps1 with or without an Mps1-6xFlag (wt or interface mutants) rescue allele. Introduction of Mps1^hm^-6xFlag cannot rescue a checkpoint deficiency, Mps1^bp^-6xFlag only partially stabilizes Pds1 compared to the wildtype. E: Representative micrographs of anaphase cells expressing Cnn1-mNeonGreen and Nuf2-mCherry with either Mps1-6xFlag or Mps1^hm^-6xFlag as their sole source of Mps1. Signals for Cnn1-mNeonGreen intensify at kinetochores during anaphase. However to a lesser extend if Mps1^hm^ is expressed. F: Quantification of time-lapse microscopy using the strains from E. Mean and SEM of all replicates are indicated as continuous lines with borders, the means of three replicates are indicated as dotted lines.

We used these alleles to assess if Mps1 interface mutants behave as “pure” checkpoint mutants, which are expected to be fully viable in absence of conditions that perturb spindle function. For instance, a checkpoint defective *mad1Δ* strain was fully viable until addition of the spindle poison Benomyl (Figure 3C). We therefore assessed the viability of Mps1 truncations (Δ111–180 and Δ144–173) and of the new point mutants we have identified. Interestingly, the truncations and the hm mutant grew poorly in the presence of NAA. Only the bp mutant had no detectable growth defect relative to the wild-type control (Figure 3C). The combination mutant Mps1^hm+bp^ had the strongest growth defect and was nearly inviable. In the presence of benomyl, as expected, none of the Mps1 interface mutants was able to confer viability. Chromosome mis-segregation and growth defects are also observed with Nuf2 CH-domain mutants that impair Mps1 kinetochore binding (Parnell et al., personal communication). These observations indicate that in addition to a requirement in SPB duplication and SAC signaling, kinetochore-localized Mps1 contributes to cell fitness under normal growth conditions, likely by supporting chromosome biorientation.

### Mps1 interface mutants are defective in SAC signalling

To better characterize the phenotype of Mps1 interface mutant, we analyzed their cell cycle progression under different conditions. To this end, we synchronized yeast cells with alpha-factor, depleted the endogenous Mps1-AID with NAA and released cells into regular medium or medium containing nocodazole to create unattached kinetochores (Figure 4A). Monitoring Pds1/Securin levels showed that Mps1-AID cells lacking an Mps1 rescue allele completed the cell cycle with similar timing as cells expressing wild-type Mps1 (Figure 4B). The Mps1 protein levels were cell-cycle regulated, as noted before^31^, with Mps1 protein declining in anaphase, slightly delayed relative to the decline in Pds1 level (Figure 4B). In the presence of Nocodazole, cells lacking an Mps1 rescue allele showed unaltered Pds1 degradation kinetics, while cells expressing wild-type Mps1 fully stabilized Pds1, indicating a functional SAC (Figure 4C). Along with Pds1, also the levels of Mps1 failed to decline in the presence of nocodazole, consistent with the notion that Mps1 itself is a substrate of the APC/C^31^. This experimental setting allowed us to test the checkpoint activity of the Mps1 interface mutants in the presence of Nocodazole (Figure 4D). We found that both Mps1^hm^ and Mps1^bp^ failed to stabilize Pds1 in the presence of nocodazole, indicating a defective mitotic checkpoint. While the Pds1 degradation kinetics of the hm mutant resembled the no-rescue situation, the bp mutant partially delayed Pds1 degradation. Interestingly, despite an active APC/C, Mps1^hm^ and Mps1^bp^ levels remained stable over time. This indicates that APC/C activation alone is not sufficient to degrade Mps1 under these conditions (in the absence of a spindle) (Figure 4 D). These results establish that Mps1 kinetochore binding is required for the SAC and that the observed SAC signal strength correlates with the residual Ndc80-binding activity of the different Mps1 interface mutants.

As the budding yeast SAC is not essential under unperturbed conditions, the SAC defect of the Mps1 interface mutants cannot explain their observed growth defects (Figure 3C). In addition, the mutants are fully proficient in SPB duplication (Supporting Figure 4B, C). To understand which additional defects may be caused by removing Mps1 from the kinetochore, we fluorescently labelled additional outer kinetochore components and analyzed their dynamic localization in the presence of either Mps1^wt^ or Mps1^hm^ by live cell imaging. The CENP-T homolog Cnn1 displays kinetochore localization at low levels during prometaphase and at increased levels as cell progress into anaphase. This increase has been suggested to depend on multiple mitotic kinases, including Mps1^32^. Indeed, we found a reduced localization of Cnn1-mNG to kinetochores in the Mps1^hm^ mutant, particularly in anaphase (Figure 4E-F). Dam1 is an established Mps1 substrate and its phosphorylation has been suggested to be involved in generating end-on attachments^33^. However, we did not observe a significant difference in the kinetochore localization of Dam1-mNeonGreen between Mps1^wt^ and Mps1^hm^ conditions (Supporting Figure 4D and E). Taken together, these observations suggest that the absence of Mps1 perturbs proper outer kinetochore architecture, which may explain the observed growth phenotypes.

### The effect of Mps1 autophosphorylation of the Ndc80c binding interface

Having verified the crucial binding interface between Mps1 and Ndc80c in cells, we set out to analyze how this interaction may be regulated. Mps1 autophosphorylation was shown to induce its release from purified kinetochore particles^25^. To identify autophosphorylation sites we incubated recombinant Mps1-StrepII with ATP in vitro and used mass spectrometry to map the relevant sites. Interestingly, out of a total of 17 detected Mps1 phosphorylation sites, six mapped in the immediate vicinity of the Mps1-Ndc80c binding interface described here (Figure 5A-C). While some of these phosphorylated residues were already annotated in the Saccharomyces Genome Database (SGD), we additionally mapped the novel phosphorylation sites Mps1:S159, Mps1:T162, and Mps1:T173. The triple cluster of phosphorylated residues Mps1:S159, Mps1:T162, and Mps1:T163 is located very close to the crucial residues Mps1:F156 and Mps1:L165, both mutated in the hm mutant (Figure 5A-C). The cluster Mps1:T173-Mps1:T175 is located very close to the invariant Mps1:R170 residue in the binding motif, while Mps1:T145 is located in the helical part of the interface. We mutated combinations of these phosphorylation sites to either alanine or aspartic acid and evaluated the phenotypes in growth assays in the Mps1-AID system. In contrast to interface mutants such as Mps1^hm^, neither of the tested phosphorylation mutants caused a significant growth defect on rich media (Figure 5D). In the presence of benomyl, however, the phosphomimetic triple mutation Mps1:S159D, Mps1:T162D, Mps1:T163D (Mps1^3D^) was inviable, while the corresponding phospho-null mutation permitted growth. We therefore focused on this triple cluster and analyzed its effect on dynamic Mps1 kinetochore localization by live cell imaging. We found that the Mps1^3D^ mutant significantly decreased Mps1 kinetochore association in prometaphase, but unlike hm or bp mutant, it did not abolish it completely (Figure 5E-G). The Mps1^3A^ mutant, on the other hand, displayed a slightly increased intensity at kinetochores during prometaphase, but was efficiently removed upon anaphase onset and did not cause a metaphase delay. Thus, phosphorylation at key residues in the Ndc80c binding interface has an effect on Mps1 kinetochore localization, but even if phosphorylation is prevented in the 3A mutant, Mps1 is efficiently removed from the kinetochore, suggesting that autophosphorylation alone is insufficient to explain the dynamic behavior of Mps1.

**Figure 5:**
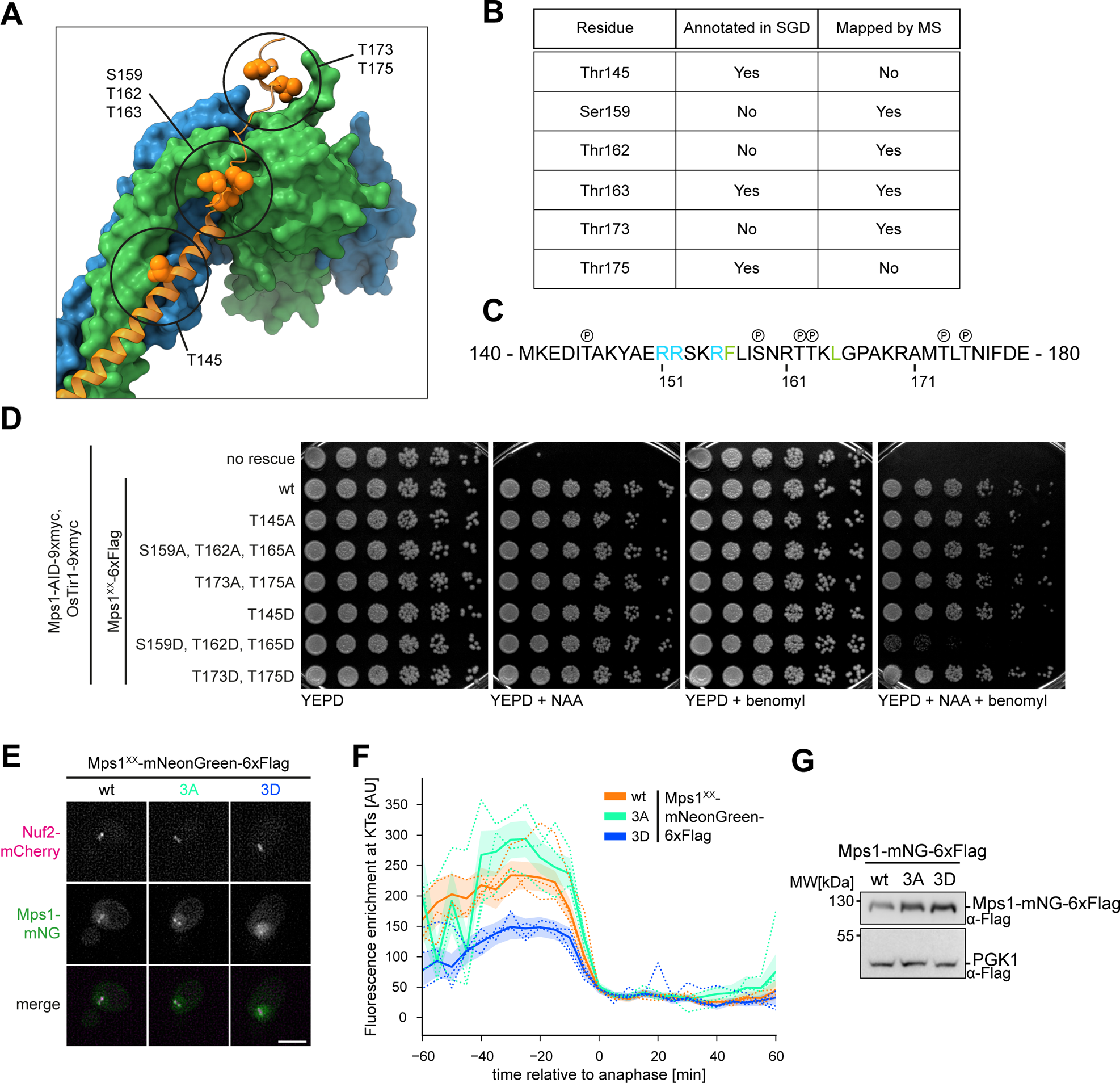
Mps1 phosphorylation in the interface region reduces kinetochore association. A: Annotation of Mps1 autophosphorylation sites within the Mps1:Ndc80c interface. Phosphorylation sites were clustered in three clusters. Cluster 1: T145; cluster 2.: S159, T162 + T163; cluster 3: T173 + T175 B: Summarizing table of Mps1 phosphorylation sites in the Mps1:Ndc80c interface. Sites were annotated in the *Saccharomyces* genome database (SGD), found in an global phosphorylation site screen, or mapped by an *in vitro* phosphorylation mass-spectrometry experiment performed in the context of this study. C: Primary sequence of Mps1^140–180^ with indications for phosphorylation sites (circled P). C: Serial dilution assay analyzing growth phenotypes of strains expressing Mps1 mutants where phosphorylation sites in the interaction interface are mutated to alanine or aspartate. Cells were spotted on YEPD, YEPD + 1 mM NAA, YEPD + 20 μg/ml benomyl and YEPD + 1 mM NAA + 20 μg/ml benomyl. Mutation of cluster 2 to aspartate results in hypersensitivity to benomyl. D: Representative micrographs of metaphase cells expressing Nuf2-mCherry and Mps1-mNeonGreen-6xFlag (wt, 3A or 3D). Mps1^3D^-mNeonGreen-6xFlag shows lower fluorescence intensities colocalizing with the signals for Nuf2-mCherry and higher signals in the nucleus compared to the wildtype or the 3A mutant. Scalebar: 5 um. E: Quantification of the kinetochore enrichment of Mps1-mNeonGreen-6xFlag (wt, 3A or 3D). Mean and SEM of all replicates are indicated as continuous lines with borders, the means of three replicates are indicated as dotted lines. F: Western blot analysis of the expression levels of Mps1-mNeonGreen-6xFlag (wt, 3A or 3D) in the strains used in D and E.

### Dam1-dependent displacement regulates Mps1 binding to Ndc80c

Given that neither microtubule-binding nor autophosphorylation may explain the dynamic regulation of the Mps1-Ndc80c binding interface, we searched for additional mechanisms controlling Mps1 kinetochore localization. The end product of the error correction pathway is a fully developed kinetochore-microtubule interface formed between Dam1 and Ndc80 complexes^34,35^. As long as attachments do not sustain tension, Ipl1 activity prevents this engagement using multiple mechanisms including downregulation of Dam1c oligomerization and direct inhibition of Ndc80c-Dam1c interactions. A recent crystallographic study has determined the binding site of the C-terminal Dam1 peptide (Dam1-C) on the Ndc80 complex^36^. Intriguingly, superposition of the Ndc80c-Dam1-C structure with the Mps1 binding interface described here, reveals substantial overlap and a similar binding mode (Figure 6A). The overlap is most obvious at the top of the Mps1 helical segment (where the AF2 prediction has a high confidence) and in the disordered segments that bind across the CH domains. The binding modalities of Mps1-N and Dam1-C peptides in this region are very similar. For example, Mps1:R170 and Dam1:R299 both establish a crucial ionic interaction with Ndc80:D295. The extensive overlap predicts that a single Ndc80 complex may be either bound to Mps1 or to Dam1-C, but not to both partners simultaneously. To test this prediction, we performed MBP-Mps1 pull-down assays with Ndc80c-Flag, but this time spiked increasing amounts of recombinant Dam1 complex into the Sf9 cell lysate. Consistent with competition for Ndc80c binding, we found that increasing concentrations of Dam1c prevented Mps1-Ndc80c interactions in a dose-dependent manner (Figure 6B). Inclusion of 1 μM Dam1c already substantially reduced binding in this setting, while 10 μM nearly completely prevented it.

**Figure 6:**
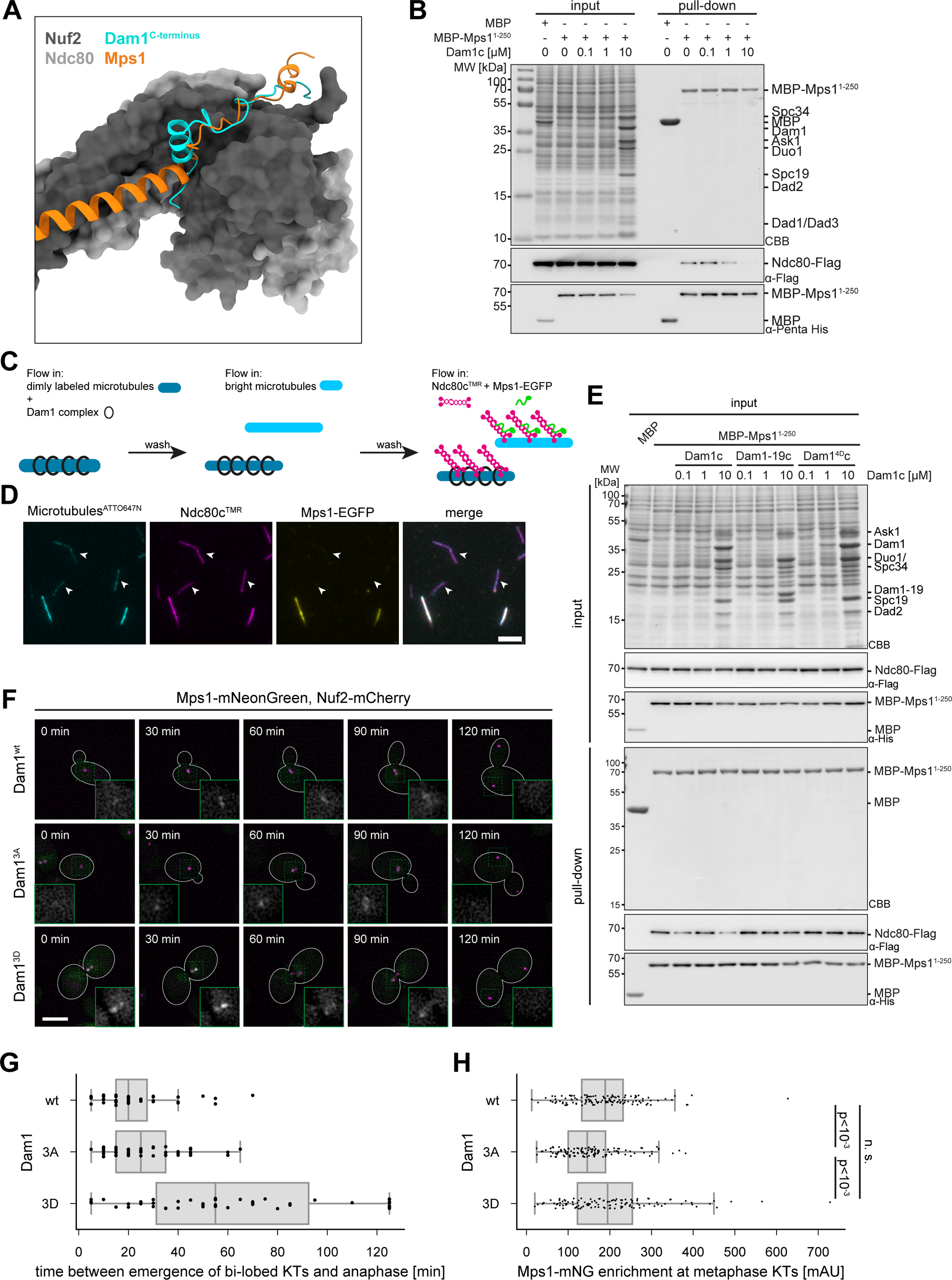
Mps1 competes with Dam1 for the interaction with the Ndc80c. A: Superposition of our Alphafold2 prediction with a crystal structure of the Ndc80:Nuf2:Dam1^C-terminus^ oligomer (PDB: 8G0Q). Mps1 and Dam1 show significant similarities concerning the mode of binding to the Ndc80c. B: Pull-down experiment using MBP-Mps1^1–250^ as bait and lysates from SF9 cells expressing the Ndc80 complex (Ndc80-Flag) spiked with recombinant Dam1c to the indicated concentrations. Increasing amounts of Dam1c in the lysate correlate with lesser amounts of Ndc80c being pulled by MBP-Mps1^1–250^. C: Schematic for a TIRF microscopy experiment to test the ability of Dam1c-decorated microtubules to recruit Ndc80c and Mps1-EGFP. D: Representative fluorescent TIRF micrographs of the experiment described in C. Mps1-EGFP colocalizes with Ndc80c-TMR on the bright microtubules but on the dimmer microtubules that were predecorated with Dam1c. Scale bar: 5 μm. Arrowheads point to dimly labeled microtubules. E: Pull-down experiment using MBP-Mps1^1–250^ as bait and lysates from SF9 cells expressing the Ndc80 complex (Ndc80-Flag) spiked with recombinant Dam1c (wt, Dam1-19 or Dam1^4D^) to the indicated concentrations. Neither Dam1-19 (lacking the C-terminus) nor Dam1^4D^ (containing mutations mimicking Ipl1 phoshphorylation of the Dam1 C-terminus) impair binding between MBP-Mps1^1–250^ and Ndc80c. F: Representative micrographs of a timelapse microscopy of cells expressing Mps1-mNeonGreen, Nuf2-mCherry in the background of Dam1 wt, 3A (S257A/S265A/S292A) or 3D (S257D/S265D/S292D). Cells expressing Dam1^3D^ show a phenotype where kinetochores are bi-lobed with over 1 μm inter-kinetochore distance but not segregated for prolonged time. Insets show the fluorescence intensities for Mps1-mNeonGreen at kinetochores. Time stamps indicate time since begin of imaging, the shown cells entered anaphase after 110 min. Scalebar: 5 μm. G: Quantification of mitotic progression of from F, measured as the time between the emergence of bi-loped kinetochore clusters and the segregation in anaphase. Cells expressing Dam1^3D^ show a delay in anaphase progression. H: Quantification of Mps1-mNeonGreen signal intensity enrichment of at metaphase kinetochores. Cells expressing Dam1^3A^ show a significant reduction in Mps1 levels. Statistical analysis: ANOVA one-way test with Dunn’s post-hoc analysis; n: over 100 cells from three independent experiments.

To visualize the relationship between Ndc80c:Dam1c and Ndc80c:Mps1 complexes more directly, we devised an experiment based on total internal reflection fluorescence (TIRF) microscopy (Figure 6C). We first decorated taxol-stabilized microtubules with recombinant Dam1 complex and immobilized them in a microscopy flow cell. We then flushed in undecorated microtubules, labeled with a higher percentage of Atto647-Tubulin to distinguish them as bright microtubules from the Dam1c-decorated, dimly labeled microtubules. Finally, an equimolar mixture of TMR-labeled Ndc80 complex and purified recombinant Mps1-EGFP was flown into the same cell. We found that Ndc80c-TMR associated with both dim (i.e. Dam1-decorated) and bright microtubules. Strikingly, Mps1-EGFP was exclusively found on bright microtubules and was absent from microtubules that were decorated by Dam1c-Ndc80c complexes (Figure 6D). We conclude that Dam1-C and Mps1 occupy the same binding site on Ndc80c and that the Dam1 complex can displace Mps1 from this interaction interface.

The C-terminus of Dam1 is a key substrate for Ipl1 during the error correction process^37^. Phosphorylation at the three residues Dam1:S257, Dam1:S265, and Dam1:S292 inhibits the interaction with Ndc80c, preventing the premature stabilization of wrongful attachments. If the Dam1-C terminus indeed competes with Mps1 for Ndc80c binding, mutations mimicking Ipl1 phosphorylation or a complete lack of the C-terminus should avoid this competition. We first tested this prediction biochemically by repeating the Ndc80c-Mps1 pull-down experiment in the presence of recombinant Dam1-19 (Q205stop mutation) or phospho-mimetic Dam1^4D^ complex (containing an S20D mutation in the Dam1 N-terminus in addition to S257D, S265D and S292D in the C-terminus). Confirming the prediction, both mutant complexes did not reduce the amount of Ndc80c bound to Mps1, even at high concentrations (Figure 6E).

We then investigated the effect of phosphorylation mutants in the Dam1 C-terminus on the recruitment of Mps1 to kinetochores in cells (Figure 6F). Interestingly, a large fraction of cells expressing Dam1^3D^ (S257D, S265D and S292D) showed a delay in metaphase (Figure 6G) and these cells were characterized by high levels of bi-lobed Mps1 signals on short spindles and a large bud size. Jin and Wang (2013) have previously shown that the Dam1^3D^ allele fully supports end-on attachments, but is specifically defective in SAC silencing^38^. Our observations here provide a mechanistic explanation for the silencing defect, because these cells fail to efficiently remove Mps1 from the kinetochore. The corresponding Dam1^3A^ allele, by contrast, resulted in decreased Mps1 accumulation at metaphase kinetochores (Figure 6H).

### Displacement of Mps1 from kinetochores occurs transiently in cells

If competition with the Dam1 C-terminus is a key mechanism to displace Mps1 from the kinetochore, the kinase should be able to re-associate under conditions in which Dam1c’s association with the kinetochore is weak. Efficient kinetochore recruitment of the Dam1 complex depends on Cdc28-dependent phosphorylation^39^ and the complex is dephosphorylated in anaphase by Cdc14^40^. To investigate Mps1 kinetochore association under conditions of low Cdk activity, we prevented the APC/C-mediated degradation of Mps1 in anaphase by mutating three candidate D-boxes in the context of the Mps1-mNeonGreen fusion protein. Confirming previous results, the mutation of the D-box at position 356 displayed the strongest effect and increased the steady-state protein level in log-phase cells^31^. Strikingly, the Mps1:Deg356 mutant re-accumulated in anaphase and became enriched again at kinetochores about 20 minutes after anaphase onset (Figure 7 A and B). This result indicates that the binding site for Mps1 at the kinetochore becomes available again as cells exit from mitosis. Furthermore, it indicates that the APC/C-dependent degradation of Mps1 is not required to induce its loss from the kinetochore, but it rather prevents its premature re-accumulation.

**Figure 7:**
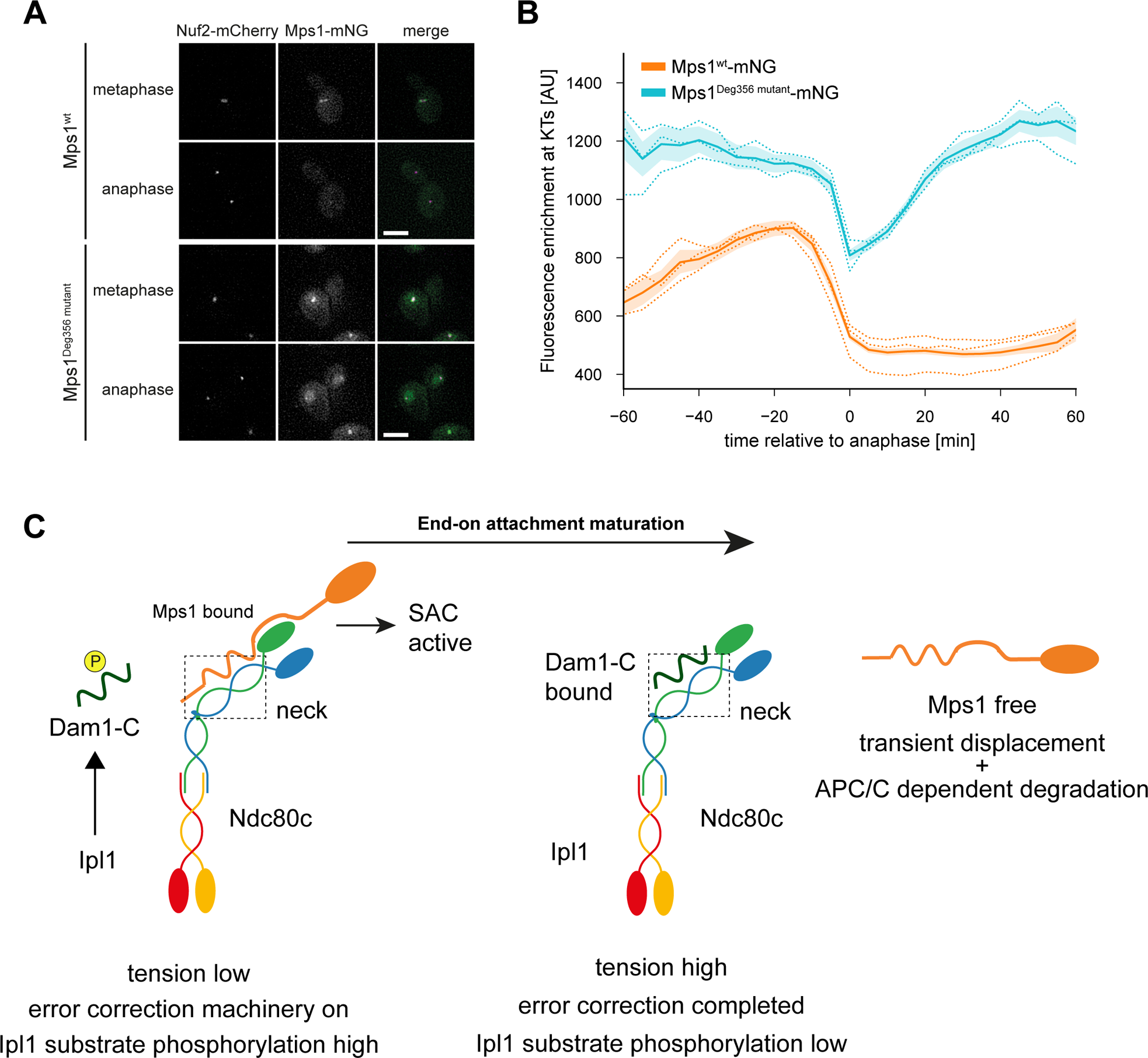
Mps1 displacement from kinetochores occurs transiently at the beginning of anaphase. A: Representative micrographs of cells in meta- and anaphase expressing Nuf2-mCherry and Mps1^wt^-mNeonGreen-6xFlag or Mps1^Deg356^ ^mutant^-mNeonGreen-6xFlag. The Deg356 mutant retains high fluorescence intensity levels in the nucleus after chromosome segregation, the fluorescence levels at kinetochores are drastically decreased. The mutant becomes enriched again at kinetochores in late anaphase. Scale bar: 5 μm. B: Quantification of fluorescence intensities Mps1-mNeonGreen-6xFlag (wt or Deg356 mutant) at the kinetochore relative to anaphase onset. Mean and SEM of all replicates are indicated as continuous lines with borders, the means of three replicates are indicated as dotted lines. C: Schematic for a model on how end-on attachment maturation leads to the displacement of Mps1 from the budding yeast kinetochore. Ipl1 phosphorylation prevents Dam1 engagement with the Ndc80c at kinetochores lacking tension. This allows Mps1 to bind and initiate SAC signaling. Once correct end-on attachments are formed, Ipl1 activity decreases, binding of the Dam1 C-terminus to the Ndc80c displaces Mps1 from the kinetochore. The SAC is silenced and Mps1 is degraded by the APC/C in anaphase which in turn prevents its premature re-association with kinetochores.

## Discussion

### The molecular architecture of the Mps1-Ndc80c interface

The experiments described here define the key binding interface between Mps1 and Ndc80c in budding yeast. In addition to our biochemical and phenotypic analysis, the crystal structure of the crucial Mps1 peptide bound to an engineered Ndc80 dwarf complex corroborates the key features of the binding mechanism (Zahm and Harrison, personal communication). The important contribution of the Nuf2 CH domain to the Mps1 recruitment mechanism is described in a related study (Parnell et al, personal communication). While we cannot fully exclude that other parts of Ndc80c or the KMN network might contribute to Mps1 binding, we note that the specificity and phenotypic severity of the described point mutants strongly argues that the described site constitutes the key binding interface. Further characterization of the binding mode with full-length proteins may offer additional insights, possibly also regarding the orientation of the Mps1 kinase domain relative to other KMN subunits. The Mps1 kinetochore recruitment mechanism presented here is fully compatible with simultaneous microtubule-binding by Ndc80c. Indeed, we show that recombinant Ndc80c recruits Mps1 to microtubules in TIRF experiments in vitro. This is in line with the demonstration of co-localization between Mps1 and native yeast kinetochores bound to microtubules^41^. Our work on the mechanism of Mps1 binding to Ndc80c rationalizes these findings.

Our findings exclude that direct binding competition between Mps1 and microtubules for Ndc80c contributes significantly to checkpoint silencing, at least in budding yeast. In this organism, Ndc80c-mediated microtubule-binding is required both for initial lateral kinetochore-microtubule attachments and for end-on attachments^42^. If initial lateral attachments prevented Mps1-binding to Ndc80c, it would be difficult to sustain the SAC during prometaphase. We note that apart from the Dam1-dependent displacement mechanism described below, the Mps1 binding site at the Ndc80c neck also offers the potential to respond to tension more directly. Microtubule-induced tension may induce conformational changes in Ndc80c, for instance by shifting the relative orientation between CH domains in the head and the coiled-coil in the shaft^43^. Given the position of the binding site, we speculate that the affinity of Mps1 for Ndc80 could be sensitive to these conformational changes. Future biophysical experiments will have to explore this possibility.

### Transient displacement and degradation regulate dynamic Mps1 association with its kinetochore receptor

A particularly revealing aspect of the binding interface identified here is its overlap with the Dam1c interaction site on Ndc80c. The assembly of a Dam1 ring complex is a hallmark of kinetochore end-on attachments in budding yeast, whereas the complex is not required for initial lateral attachments^42^. The maturation of the Dam1c:Ndc80 interface is a key element of control during error correction with Ipl1 preventing wrongful end-on attachments by phosphorylating the C-terminus of Dam1^44^. We propose that rather than a simple competition for microtubule binding, the gradual replacement of Mps1 with Dam1c during attachment maturation makes the availability of the Mps1 kinetochore binding site dependent on reaching a specific attachment configuration (Figure 7C). Our experiments with the Mps1 degron mutant reveal that the displacement mechanism only operates transiently. In late anaphase, the affinity between Dam1c and Ndc80c may decrease again, possibly due to Cdc14-dependent dephosphorylation of Ask1^40^. In principle, this would make the binding site for Mps1 available again, were it not for the APC/C-dependent degradation of Mps1. This finding highlights a fine balance between overall Mps1 protein levels and the availability of a kinetochore binding site.

A hallmark of Mps1 kinetochore recruitment in different systems is its sensitivity to Mps1 kinase activity, i.e. inhibition of Mps1 kinase activity typically promotes its kinetochore localization. Consistent with this, our results show that key Mps1 autophosphorylation sites are located at the Ndc80c binding interface and that phospho-mimetic mutations reduce the Mps1 level at the kinetochore. Our analysis of the non-phosphorylatable Mps1^3A^ mutant argues that autophosphorylation at these sites is not strictly required for the kinetochore release of Mps1. It remains possible, however, that autophosphorylation could facilitate the Dam1-dependent displacement mechanism. A simultaneous increase in Mps1 autophosphorylation and decrease in Ipl1 phosphorylation upon correct end-on attachment may tip the balance, causing Dam1c recruitment and Mps1 displacement.

The molecular details of the Mps1 binding mechanism to Ndc80c do not seem to be fully conserved between yeast and humans. The human Mps1 N-terminus contains a TPR domain and an N-terminal extension, which do not seem to have a structural equivalent in yeast Mps1. We propose, however, that the concept of Mps1 displacement during attachment maturation may also apply to human kinetochores and could constitute a general molecular strategy to regulate Mps1.

### Insights into the overall coordination between error correction and SAC

The findings described here illuminate novel aspects of the relationship between the tension-sensitive error correction machinery and the spindle assembly checkpoint and they further specify the role of Ipl1/Aurora B in the SAC in budding yeast. The Dam1 C-terminus is regulated by Ipl1 during error correction, placing the Ndc80c:Mps1 association under Ipl1 control, at least indirectly (Figure 7C). As long as Ipl1 is active (Ipl1 substrate phosphorylation high), Dam1-C is prevented from competing away Mps1. This relationship can explain a number of observations regarding differential requirements for Ipl1 activity in the SAC. Previous studies have shown, for example, that Ipl1 activity is required for the metaphase arrest induced by Mps1 overexpression, i.e. pGal-Mps1 cells cannot maintain a metaphase arrest in *ipl1* mutants^45^. Our findings argue that in this situation, dephosphorylation of the Dam1-C terminus at the restrictive temperature should make it difficult for Mps1 to bind Ndc80c, weakening the SAC. On the other hand, Ipl1 activity is not required for the SAC response following microtubule depolymerization by nocodazole^46^. Given that the kinetochore association of Dam1 is largely microtubule-dependent^47^, we suggest that in this situation Mps1 does not have to compete against Dam1c for Ndc80c binding, making Ipl1 activity unnecessary for a SAC response. Furthermore, *ipl1* temperature-sensitive mutants fail to activate the SAC at the restrictive temperature despite extensive attachment errors^48^. A key problem that *ipl1* mutants face in this situation may be the difficulty to compete away Dam1 from Ndc80c for Mps1 binding. In line with this idea, *ipl1 dam1* double mutants have a functional SAC, even though *dam1* mutants typically do not create unattached kinetochores^49^. Therefore, in budding yeast the error correction machinery has an essential and dominating role in regulating kinetochore-microtubule attachments and it also controls key upstream events of the SAC, such as regulated Mps1 recruitment and release.

## Methods

### Yeast strains

| Name | Genotype | source | Figure |
| --- | --- | --- | --- |
| RPY115 | Mat a, Ade2, leu2-3,112::Mps1-mNeonGreen-6xFlag::Leu2, his3D200, ura3-52::pTEF1_OsTir1_tADH1, lys2-801, Mps1-AID-9xmyc::KanMX6; Nuf2-mCherry::His3MX6 | This study | 3A+B, 4D+E 7A+B |
| RPY116 | Mat a, Ade2, leu2-3,112::Mps1(F156D, L165A)-mNeonGreen-6xFlag::Leu2, his3D200, ura3-52::pTEF1_OsTir1_tADH1, lys2-801, Mps1-AID-9xmyc::KanMX6; Nuf2-mCherry::His3MX6 | This study | 3A+B |
| RPY117 | Mat a, Ade2, leu2-3,112::Mps1(R151D, R152D, R155D)-mNeonGreen-6xFlag::Leu2, his3D200, ura3-52::pTEF1_OsTir1_tADH1, lys2-801, Mps1-AID-9xmyc::KanMX6; Nuf2-mCherry::His3MX6 | This study | 3A+B |
| RPY121 | Mat a, Ade2, leu2-3,112::Mps1(R151D, R152D, R155D, F156D, L165A)-mNeonGreen-6xFlag::Leu2, his3D200, ura3-52::pTEF1_OsTir1_tADH1, lys2-801, Mps1-AID-9xmyc::KanMX6; Nuf2-mCherry::His3MX6 | This study | 3A+B |
| RPY80a | Mat a, Ade2, leu2-3,122, his3D200, ura3-52::pTEF1_OsTir1_tADH1, lys2-801, Mps1-AID-9xmyc::KanMX6 | This study | 3C, S3D+E |
| RPY82a | Mat a, Ade2, leu2-3,112::Mps1-6xFlag::Leu2, his3D200, ura3-52::pTEF1_OsTir1_tADH1, lys2-801, Mps1-AID-9xmyc::KanMX6 | This study | 3C, S3D+E |
| RPY91 | Mat a, Ade2, leu2-3,122::Mps1(del 144-173)-6xFlag::Leu2, his3D200, ura3-52::pTEF1_OsTir1_tADH1, lys2-801, Mps1-AID-9xmyc::KanMX6 | This study | 3C |
| RPY92 | Mat a, Ade2, leu2-3,122::Mps1(del 111-180)-6xFlag::Leu2, his3D200, ura3-52::pTEF1_OsTir1_tADH1, lys2-801, Mps1-AID-9xmyc::KanMX6 | This study | 3C |
| RPY93 | Mat a, Ade2, leu2-3,122::Mps1(F165D, L165A)-6xFlag::Leu2, his3D200, ura3-52::pTEF1_OsTir1_tADH1, lys2-801, Mps1-AID-9xmyc::KanMX6 | This study | 3C |
| RPY94 | Mat a, Ade2, leu2-3,122::Mps1(R151D, R152D, R155D)-6xFlag::Leu2, his3D200, ura3-52::pTEF1_OsTir1_tADH1, lys2-801, Mps1-AID-9xmyc::KanMX6 | This study | 3C |
| RPY105 | Mat a, Ade2, leu2-3,122::Mps1(F165D, L165A)-6xFlag::Leu2, his3D200, ura3-52::pTEF1_OsTir1_tADH1, lys2-801, Mps1-AID-9xmyc::KanMX6, Pds1-3xHA::His3MX6 | This study | 3C |
| RPY106 | Mat a, Ade2, leu2-3,122::Mps1(R151D, R152D, R155D)-6xFlag::Leu2, his3D200, ura3-52::pTEF1_OsTir1_tADH1, lys2-801, Mps1-AID-9xmyc::KanMX6, Pds1-3xHA::His3MX6 | This study | 3C |
| RPY109 | Mat a, Ade2, leu2-3,122::Mps1(F165D, L165A)-6xFlag::Leu2, his3D200, ura3-52::pTEF1_OsTir1_tADH1, lys2-801, Mps1-AID-9xmyc::KanMX6, Spc42-mCherry::His3MX6 | This study | 3C |
| RPY87 | Mat a, Ade2, leu2-3,122, his3D200, ura3-52::pTEF1_OsTir1_tADH1, lys2-801, Mps1-AID-9xmyc::KanMX6, Pdc1-3xHA::HIS3MX6 | This study | 4B+C+D |
| RPY88 | Mat a, Ade2, leu2-3,112::Mps1-6xFlag::Leu2, his3D200, ura3-52::pTEF1_OsTir1_tADH1, lys2-801, Mps1-AID-9xmyc::KanMX6, Pdc1-3xHA::HIS3MX6 | This study | 4B+C+D |
| RPY162 | Mat a, Ade2, leu2-3,112::Mps1(F156Dm L165A)-6xFlag::Leu2, his3D200, ura3-52::pTEF1_OsTir1_tADH1, lys2-801, Mps1-AID-9xmyc::KanMX6; Nuf2-mCherry::His3MX6, Cnn1-mNeonGreen::hphNT1 | This study | 4E+F |
| RPY163 | Mat a, Ade2, leu2-3,112::Mps1-6xFlag::Leu2, his3D200, ura3-52::pTEF1_OsTir1_tADH1, lys2-801, Mps1-AID-9xmyc::KanMX6; Nuf2-mCherry::His3MX6, Cnn1-mNeonGreen::hphNT1 | This study | 4E+F |
| RPY84 | Mat a, Ade2, leu2-3,122::Mps1(T145A)-6xFlag, his3D200, ura3-52::pTEF1_OsTir1_tADH1, lys2-801, Mps1-AID-9xmyc::KanMX6 | This study | 5C |
| RPY85 | Mat a, Ade2, leu2-3,122::Mps1(S159A, T162A, T163A)-6xFlag, his3D200, ura3-52::pTEF1_OsTir1_tADH1, lys2-801, Mps1-AID-9xmyc::KanMX6 | This study | 5C |
| RPY86 | Mat a, Ade2, leu2-3,122::Mps1(T173A, T175A)-6xFlag, his3D200, ura3-52::pTEF1_OsTir1_tADH1, lys2-801, Mps1-AID-9xmyc::KanMX6 | This study | 5C |
| RPY127 | Mat a, Ade2, leu2-3,122::Mps1(T145D)-6xFlag, his3D200, ura3-52::pTEF1_OsTir1_tADH1, lys2-801, Mps1-AID-9xmyc::KanMX6 | This study | 5C |
| RPY128 | Mat a, Ade2, leu2-3,122::Mps1(S159D, T162D, T163D)-6xFlag, his3D200, ura3-52::pTEF1_OsTir1_tADH1, lys2-801, Mps1-AID-9xmyc::KanMX6 | This study | 5C |
| RPY129 | Mat a, Ade2, leu2-3,122::Mps1(T173D, T175D)-6xFlag, his3D200, ura3-52::pTEF1_OsTir1_tADH1, lys2-801, Mps1-AID-9xmyc::KanMX6 | This study | 5C |
| RPY145 | Mat a, Ade2, leu2-3,112::Mps1(S159A, T162A, T163A)-mNeonGreen-6xFlag::Leu2, his3D200, ura3-52::pTEF1_OsTir1_tADH1, lys2-801, Mps1-AID-9xmyc::KanMX6; Nuf2-mCherry::His3MX6 | This study | 5D+E |
| RPY147 | Mat a, Ade2, leu2-3,112::Mps1(S159D, T162D, T163D)-mNeonGreen-6xFlag::Leu2, his3D200, ura3-52::pTEF1_OsTir1_tADH1, lys2-801, Mps1-AID-9xmyc::KanMX6; Nuf2-mCherry::His3MX6 | This study | 5D+E |
| RPY72 | Mat a, ade2-1, leu2-3,112, his3D200, ura3-52::pTEF1-OsTIR1_tADH1::URA3, Lys2 | This study | S3D+E |
| RPY79 | Mat a, ade2-1, leu2-3,112::Mps1-6xFlag::Leu2, his3D200, ura3-52, Lys2, Mps1-AID-9xmyc::KanMX6 | This study | S3D+E |
| RPY89 | Mat a, Ade2, leu2-3,122, his3D200, ura3-52::pTEF1_OsTir1_tADH1, lys2-801, Mps1-AID-9xmyc::KanMX6, Spc42-mCherry::HIS3MX6 | This study | S4B+C |
| RPY90 | Mat a, Ade2, leu2-3,112::Mps1-6xFlag::Leu2, his3D200, ura3-52::pTEF1_OsTir1_tADH1, lys2-801, Mps1-AID-9xmyc::KanMX6, Spc42-mCherry::HIS3MX6 | This study | S4B+C |
| RPY110 | Mat a, Ade2, leu2-3,122::Mps1(R151D, R152D, R155D)-6xFlag::Leu2, his3D200, ura3-52::pTEF1_OsTir1_tADH1, lys2-801, Mps1-AID-9xmyc::KanMX6, Spc42-mCherry::His3MX6 | This study | S4B+C |
| RPY152 | Mat a, Ade2, leu2-3,112::Mps1(F156Dm L165A)-6xFlag::Leu2, his3D200, ura3-52::pTEF1_OsTir1_tADH1, lys2-801, Mps1-AID-9xmyc::KanMX6; Nuf2-mCherry::His3MX6, Dam1-mNeonGreen::hphNT1 | This study | S4D+E |
| RPY153 | Mat a, Ade2, leu2-3,112::Mps1-6xFlag::Leu2, his3D200, ura3-52::pTEF1_OsTir1_tADH1, lys2-801, Mps1-AID-9xmyc::KanMX6; Nuf2-mCherry::His3MX6, Dam1-mNeonGreen::hphNT1 | This study | S4D+E |
| ZNY12 | Mat a, Ade2, leu2-3,112::Mps1(RL356/358AA)-mNeonGreen-6xFlag::Leu2, his3D200, ura3-52::pTEF1_OsTir1_tADH1, lys2-801, Mps1-AID-9xmyc::KanMX6; Nuf2-mCherry::His3MX6 | This study | 3A+B,<br>4D+E<br>7A+B |

### Plasmids

| Plasmid | Coding elements | Source | Purpose |
| --- | --- | --- | --- |
| pRP15 | pRP305_Mps1(delta144-173)-6xFlag | This study | Yeast transformation |
| pRP23 | pRS305_MPS1(T145A)-6xFlag | This study | Yeast transformation |
| pRP24 | pRS305_MPS1(S159A, T162A, T163A)-6xFlag | This study | Yeast transformation |
| pRP25 | pRS305_MPS1(T173A, T175A)-6xFlag | This study | Yeast transformation |
| pRP36 | pET-Duet_MBP-Mps1(1-250)-8xHis | This study | Protein purification |
| pRP41 | pRP305_Mps1(delta111-180)-6xFlag | This study | Yeast transformation |
| pRP45 | pRS305_Mps1-mNeonGreen-6xFlag | This study | Yeast transformation |
| pRP46 | pRP305_Mps1(hm)-6xFlag | This study | Yeast transformation |
| pRP47 | pRP305_Mps1(bp)-6xFlag | This study | Yeast transformation |
| pRP48 | pET-Duet_MBP-Mps1(1-250, hm)-8xHis | This study | Protein purification |
| pRP49 | pET-Duet_MBP-Mps1(1-250, bp)-8xHis | This study | Protein purification |
| pRP50 | pRP305_Mps1(hm+bp)-6xFlag | This study | Yeast transformation |
| pRP52 | pET-Duet_MBP-Mps1(1-250, hm+bp)-8xHis | This study | Protein purification |
| pRP53 | pRS305-MPS1(T145D)-6xFlag | This study | Yeast transformation |
| pRP53 | pRS305-MPS1(S159D, T162D, T163D)-6xFlag | This study | Yeast transformation |
| pRP55 | pRS305-MPS1(T173D, T175D)-6xFlag | This study | Yeast transformation |
| pRP57 | pRS305_Mps1-mNeonGreen(hm)-6xFlag | This study | Yeast transformation |
| pRP58 | pRS305_Mps1-mNeonGreen(bp)-6xFlag | This study | Yeast transformation |
| pRP59 | pRS305_Mps1-mNeonGreen(hm+bp)-6xFlag | This study | Yeast transformation |
| pRP63 | pFL_StrepII-Mps1(hm) | This study | Protein purification |
| pRP64 | pFL_StrepII-Mps1(bp) | This study | Protein purification |
| pRP65 | pFL_StrepII-Mps1(hm+bp) | This study | Protein purification |
|  | pET-Duet_MBP-8xHis | Musacchio lab | Protein purification |
|  | pBig1_Ndc80-Flag :Nuf2 :Spc24:Spc25 | Westermann Lab | Protein purification |
|  | pFL_Strep2-Mps1 | Westermann Lab | Protein purification |

### Protein purification

MBP-Mps1^1–250–^8xHis fragments were expressed from pET-Duet derived plasmids in BL21 *E. coli* cells. Cells containing the plasmid grown in 1 L TB media containing 100 mg/L ampicillin at 37 °C to OD600 = 0.6-0.7, expression was induced through the addition of IPTG to a final concentration of 0.2 mM. Cells were harvested after a 20 h expression period at 20 °C. *E. coli* cells were suspended in lysis buffer (500 mM NaCl, 30 mM Hepes pH = 7.5, 10 % glycerol, 30 mM imidazole, 0.5 mM TCEP, 0.01 % TritonX-100) and lysed by sonication. Insoluble components were separated through centrifugation (6,000 xg, 4 °C, 20 min). The supernatant was incubated with 1 ml NiNTA Superflow resin (Takara) overnight rotating at 4 °C. The loaded beads were washed with 100 ml lysis buffer. The bound proteins were eluted with elution buffer (500 mM NaCl, 30 mM Hepes pH = 7.5, 10 % glycerol, 400 mM imidazole, 0.5 mM TCEP, 0.01 % TritonX-100). The eluate further purified by size exclusion chromatography using a Superdex200 10/300 increase (GE healthcare; buffer: 300 mM NaCl, 5 % glycerol, 20 mM Hepes pH = 7.5, 0.5 mM TCEP). The peak fractions were pooled, concentrated, snap frozen in liquid nitrogen and stored at −80 °C.

Mps1-EGFP bearing an N-terminal Strep-tag II was expressed in *Sf9* insect cells. Insect cell pellets were resuspended in lysis buffer (20mM Na_2_HPO_4_/NaH_2_PO_4_ pH 7.5, 300 mM NaCl, 5% glycerol, 0.5% NP40 alternative, 1 mM PMSF, cOmplete protease inhibitor (Roche) and PhoSTOP phosphatase inhibitors (Roche)) and lysed by sonication. The lysate was cleared by centrifugation (21,000 xg, 30 min, 4°C), mixed with Strep-Tactin Superflow Plus resin (Qiagen) and incubated overnight at 4°C. The resin was washed extensively with wash buffer (20 mM Na_2_HPO_4_/NaH_2_PO_4_ pH 7.5, 300 mM NaCl, 5% glycerol, 0.1% Tween20) and the protein was eluted with wash buffer supplemented with 5 mM d-desthiobiotin and 0.5 mM TCEP. Once eluted, Mps-EGFP was kept on ice and used for TIRF microscopy assays within 24 h.

Ndc80c bearing a FLAG-tag on the C-terminus of the Ndc80 subunit and a Halo-tag on the C-terminus of Spc25 was expressed in *Sf9* insect cells. Insect cell pellets were resuspended in lysis buffer (20mM Na_2_HPO_4_/NaH_2_PO_4_ pH 7.5, 300 mM NaCl, 5% glycerol, 0.5% NP40 alternative, 1 mM PMSF cOmplete protease inhibitor (Roche) and PhoSTOP phosphatase inhibitors (Roche)) and lysed by sonication. After clearing by centrifugation (21,000 xg, 30 min, 4°C), the lysate was mixed with anti-FLAG M2 affinity gel (Sigma-Aldrich) and incubated overnight at 4°C. The resin was washed extensively with wash buffer (20 mM Na_2_HPO_4_/NaH_2_PO_4_ pH 7.5, 300 mM NaCl, 5% glycerol) and Ndc80c was fluorescently labeled with HaloTag TMR ligand (Promega) for 2 h at 4°C. The unbound TMR ligand was removed by extensively washing the resin with wash buffer supplemented with 0.1% Tween20. Proteins were eluted with wash buffer supplemented with 0.5 mg/mL 3xFLAG peptide, 0.1% Tween20, and 0.5 mM TCEP. Once eluted, Ndc80c-Halo^TMR^ was kept on ice and used for TIRF microscopy assays within 24 h.

Tubulin was purified from porcine brain as described before (Castoldi and Popov, 2003) and labeled with biotin-N-hydroxysuccinimide (NHS; Sigma-Aldrich) or ATTO-647N-NHS (ATTO-TEC) following standard procedures (Hirst et al., 2020).

Dam1 complexes were purified as described previously (Dudziak et al., 2021).

### Pull-down assay

*Sf9* insect cells were infected to express the Ndc80 complex (Ndc80-Flag:Nuf2:Spc24:Spc25). Cells were suspended in lysis buffer (200 mM NaCl, 20 mM Hepes pH = 7.5, 1 mM DTT, 0.01 % Tween20) supplemented with cOmplete protease inhibitor (Roche), PhoSTOP phosphatase inhibitors (Roche) and 1 mM PMSF and lysed by sonication. The lysate was cleared by centrifugation (21,000 xg, 30 min, 4 °C). 200 pmol MBP or MBP-Mps1^1–250^ was immobilized on amylose resin (New England BioLabs) equilibrated in lysis buffer. The lysate was mixed with the loaded beads and incubated at 4 °C for 1 h rotating. The beads were washed five times with lysis buffer. Proteins on the beads were eluted with SDS-sample buffer. Samples were analyzed by Coomassie stained SDS-PAGEs and western blots.

For the Dam1c competition assay, the lysate was mixed 1:1 with Dam1c for final concentration of 0.1 μM, 1 μM or 10μM.

### Co-immunoprecipitation and size exclusion chromatography

Ndc80c bearing a FLAG-tag on the C-terminus of the Ndc80 subunit and Mps1 (wt, hm or hm+bp) bearing a Strep-tag II on the N-terminus were co-expressed in *Sf9* cells. The insect cell pellets were resuspended in lysis buffer (300 mM NaCl, 20 mM Na_2_HPO_4_/NaH_2_PO_4_ pH = 7.5, 2.5 % glycerol, 0.5 % NP-40) and lysed by sonication. The lysate was cleared by centrifugation (21,000 xg, 30 min, 4 °C). The cleared lysate was mixed with M2-Flag resins and incubated rotating overnight at 4 °C. The loaded beads were washed with wash buffer (300 mM NaCl, 20 mM Na_2_HPO_4_/NaH_2_PO_4_ pH = 7.5, 2.5 % glycerol, 0.01 % Tween20). Proteins were eluted with elution buffer (300 mM NaCl, 20 mM Na_2_HPO_4_/NaH_2_PO_4_ pH = 7.5, 2.5 % glycerol, 0.01 % Tween20, 0.5 mg/ml 3xFlag peptide). The protein elution was run over a Superrose 6 3.2/300 size exclusion chromatography column equilibrated in chromatography buffer (300 mM NaCl, 20 mM Na_2_HPO_4_/NaH_2_PO_4_ pH = 7.5, 2.5 % glycerol).

### Cell culture

Yeast cells were grown in Yeast extract peptone dextrose (YEPD) media at 30 °C. Live cell imaging was done in synthetic media supplemented with essential amino acids, glucose, and if indicated 1 mM NAA. Western blot samples were generated according to standard methods (Kushnirov, 2000).

Growth phenotypes were characterized through serial dilution assays. Cells of the indicated strain were diluted to an OD = 0.4 in the first well of an 96-well plate and diluted five times 1:4 in the following wells. Equal drops of cells were transferred using a 48-pin frogger on YEPD plates, supplemented with 1 mM NAA or 20 μg/ml benomyl if indicated. The plates were incubated at 30°C for two days.

### Yeast strain construction

Transformations were performed according to standard methods (Schiestl and Gietz, 1989).

Mps1 integration constructs were generated as pRS305 derived plasmids. The Mps1 open reading frame together with 300bp of the 3’ and 5’ untranslated region was integrated into the multiple cloning site using Gibson assembly. Epitope tags (6xFlag or mNeonGreen-6xFlag) were introduced at the C-terminus of Mps1. Mps1 point mutations were integrated through site directed mutagenesis PCR. The integration constructs were linearized by cleavage with the restriction enzyme *AflII* to be integrated at the Leu2 locus in the budding yeast genome.

Additional epitope tags (Spc42-mCherry, Nuf2-mCherry, Mps1-AID-9xmyc, Pds1-3xHA, Cnn1-mNeonGreen, Dam1-mNeonGreen) were introduced at the endogenous locus of the respective protein through PCR based tagging methods ^50,51^.

### Spindle pole body duplication assay

YEPD day cultures of yeast strains (Spc42-mCherry, Mps1-AID-9xmyc, Mps1^XX^-6xFlag, OsTir-9xmyc) were inoculated from overnight cultures to an OD = 0.2 and grown for 2.5 h at 30 °C. Depletion of the endogenous Mps1 was induced by addition of NAA to 1 mM final concentration. Cells were harvested 1.5 h after NAA addition and fixed by treatment with 4 % PFA for 15 min. Images were acquired with a DeltaVision elite system (objective: Plan Apo 60x/1.42 oil DIC; camera: pco.edge 5.5 sCMOS). Large-budded cells were examined to show one or two distinct signals for Spc42-mCherry using ImageJ utilizing the YeastMate plugin^52^.

### Cell cycle progression assay

YEPD day cultures of yeast strains (Pds1-3xHA, Mps1-AID-9xmyc, Mps1^XX^-6xFlag, OsTir-9xmyc) were inoculated from overnight cultures to an OD = 0.15 and grown at 30 °C for 1 h. G1-phase arrest was induced by adding α-factor peptide to 10 μg/ml final concentration. NAA was added after 90 min to 1 mM final concentration. The α-factor was washed out 120 min after its addition by washing twice with YEPD containing 1 mM NAA and 1 mg/ml pronase and once with YEPD containing 1 mM NAA. Cells were released in YEPD containing 1 mM NAA +/− 20 μg/ml nocodazole. Samples for WB analysis were taken every 15 min for 2 h. Initiation of a second cell cycle was prevented by adding α-factor 45 min after release.

### Live cell microscopy

YEPD day cultures of yeast strains were inoculated from overnight cultures to an OD = 0.2 and grown at 30 °C for three to four hours. Cells were suspended in imaging media containing 1 mM NAA and immobilized in 15μ-slides (Ibidy) precoated with concanavalin A. The slide was mounted on a DeltaVision Elite system (objective: Super-Plan Apo 100x/1.4 oil DIC; camera: pco.edge 5.5 sCMOS) preheated to 30 °C. Timelapse images were captured every 5 min for 90 or 120 min.

Image analysis was done in ImageJ. Dividing cells were selected based on the separation of Nuf2-mCherry signals. An ImageJ macro extracted the mean intensity values at kinetochore clusters (Nuf2-mCherry foci) as well as in the nucleus (0.2 μm band around the Nuf2-mChery foci). Kinetochore enrichments were calculated by subtracting the intensity value in the nucleus from the intensity value at the kinetochore cluster. Values were plotted relative to the first anaphase frame.

### TIRF microscopy

Microtubules were assembled from 20 µM tubulin in BRB80 (80 mM PIPES, pH 6.8 with KOH, 1mM MgCl_2_, 1 mM EGTA) supplemented with 2 mM MgCl_2_, 1 mM DTT, 25% glycerol, and 1 mM GTP for 30 min at 37°C. For dim microtubules, the mixture contained 5% ATTO-647N-labeled tubulin, 25% biotinylated tubulin and 70% unlabeled tubulin, whereas for bright microtubules the mixture contained 25% ATTO-647N-labeled tubulin, 25% biotinylated tubulin and 50% unlabeled tubulin. Microtubules were stabilized with 20 µM taxol and maintained at 37°C at least 15 minutes before being diluted 1/100 in BRB80 supplemented with 1 mg/mL β-casein, 0.05% methylcellulose, 1 µM taxol, and an oxygen scavenging mix (200 μg/ml glucose-oxidase, 35 μg/ml catalase, 10 mM, and 4.5 μg/ml glucose).

Flow chambers were assembled with double-sided tape between a cleaned glass slide and a biotin-PEG functionalized coverslip as described previously (Hirst et al., 2020). Chambers were first blocked with 1% Pluronic F-127 for 30 min and then with a mixture of 1 mg/mL β-casein and 0.1 mg/mL avidin DN (Vector Laboratories) for at least 30 min. Flow chambers were equilibrated with assay buffer (BRB80 supplemented with 1 mg/mL β-casein, 0.15% methylcellulose, 10% glycerol, 0.02% Brij-35, 1 µM taxol, 200 μg/ml glucose-oxidase, 35 μg/ml catalase, 10 mM, and 4.5 μg/ml glucose), then dim microtubules were introduced and allowed to attach to the surface for 30 sec. Unbound microtubules were washed away with assay buffer and then a mixture of 1 µM Dam1c in assay buffer was flown in the chamber. Dam1c was allowed to form rings around dimly labeled microtubules for 10 minutes, and then washed away with assay buffer. Next, brightly labeled microtubules were introduced and allowed to attach to the surface for 30 seconds. The chamber was washed again with assay buffer before introducing 100 nM Mps1-EGFP and 100 nM Ndc80c-HaloTMR in assay buffer. The slide was visualized at 30°C on a Nikon Ti2 Eclipse microscope equipped with an oil immersion objective (Nikon, CFI Apochromat TIRF 100x, NA1.49) and an Andor iXon Life 888 EMCCD camera.

## Acknowledgments

The authors thank Steve Harrison and Matt Miller for communicating results prior to publication. This work was funded by the Deutsche Forschungsgemeinschaft (DFG, German Research Foundation) – SFB1430 Molecular Mechanisms of Cell State Transitions – Project-ID 424228829. Microscopy experiments were carried out with support from the Imaging Center Campus Essen Core Facility (ICCE). The DeltaVision Elite high resolution microscope was obtained through Deutsche Forschungsgemeinschaft funding (Major Research Instrumentation Programme as per Art. 91b GG, INST 20876/275-1). The wide-field TIRF microscope was obtained through Deutsche Forschungsgemeinschaft funding (Major Research Instrumentation Programme as per Art. 91b GG, INST 20876/276-1).

## Author Contributions

RP conducted and analyzed the majority of experiments. CC performed TIRF experiments. SH supported recombinant protein expression. CD performed initial characterization of mutants. AD conducted phosphorylation experiments. FK performed mass spec analysis. AM and IV performed structure predictions and sequence analysis. SW acquired funding and conceived the study. RP and SW wrote the manuscript with input from all authors.

## Conflict of Interest

The authors declare no conflict of interest.

---

**Supporting Figure 1:**
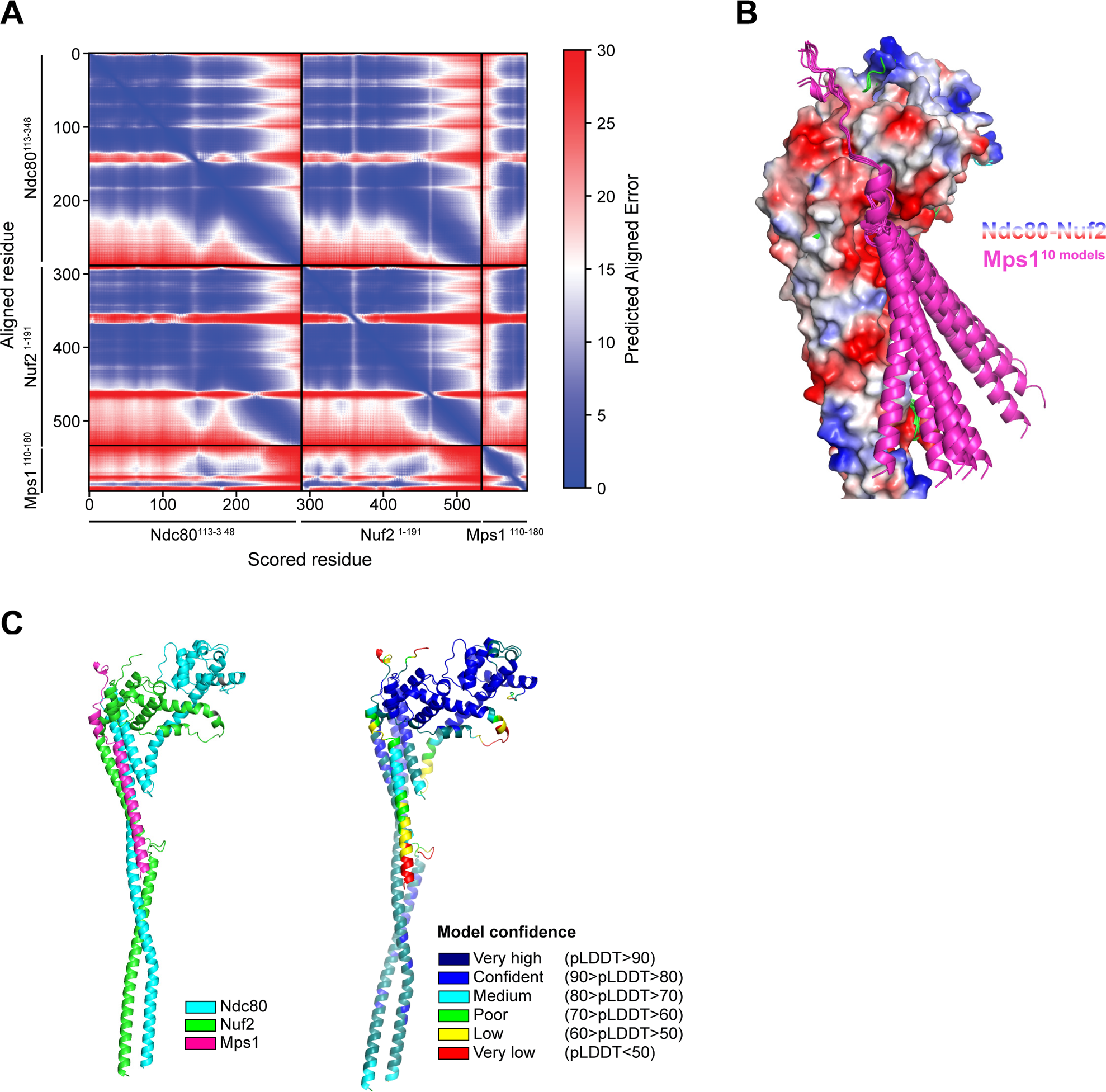
Alphafold2 prediction of the Ndc80:Nuf2:Mps1 oligomer. A: Predicted alignment error of the Alpha-fold 2 prediction of the Ndc80^113–438^: Nuf2^1–191^: Mps1^110–180^ oligomer. B: A superposition of ten Alphafold2 predictions of the Ndc80^113–438^: Nuf2^1–191^: Mps1^110–180^ oligomer. The ten structure predictions for the Mps1 segment are shown in magenta. They confidently align in the unstructured element and diverge in the helical segment. The surface of Nuf2 and Ndc80 is colored by its theoretical charge. C: Left: The Ndc80^113–438^: Nuf2^1–191^: Mps1^110–180^ oligomer structure colored by chain identity. Right: The Ndc80^113–438^: Nuf2^1–191^: Mps1^110–180^ oligomer structure colored by pLDDT model confidence. For Mps1 the model shows the highest confidence in the unstructured loop segment that binds in the neck of the CH-domains.

**Supporting Figure 3.**
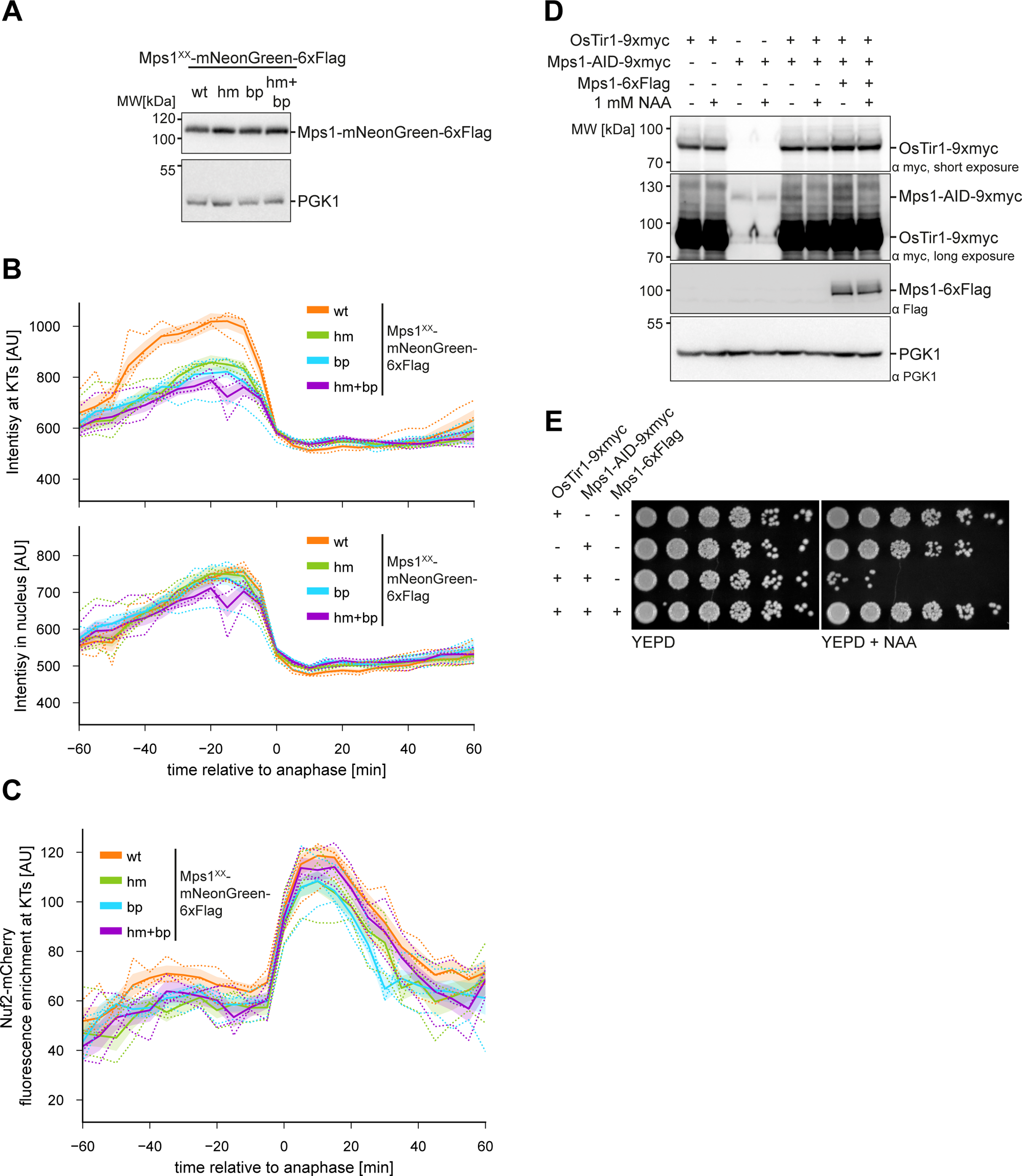
Additional characterization of live cell microscopy and Mps1-AID alleles. A: Western blot analysis showing Mps1-mNeonGreen-6xFlag (wt or interface mutants) are expressed to comparable amounts. B: Quantification of fluorescence signal intensities for Mps1-mNeonGreen-6xFlag(wt or interface mutants) at kinetochores (upper panel) or in the nuclear background (lower panel) relative to the first anaphase frame. Both wildtype and the interface mutants show an increase in prometa- and metaphase and a sharp decrease as cells enter anaphase, reflecting the cell cycle regulated protein stability of Mps1. Mean and SEM of all replicates are indicated as continuous lines with borders, the means of three replicates are indicated as dotted lines. C: Quantification of kinetochore enrichment of Nuf2-mCherry relative to the first anaphase frame. Mean and SEM of all replicates are indicated as continuous line with borders, the means of individual replicates are indicated as dotted lines. D: Western blot analysis of the depletion of the endogenous Mps1. The depletion only occurs when OsTir1 and Mps1-AID are combined in the same strain, protein levels of the exogenous Mps1-6xFlag allele without AID tag are not altered upon treatment. Samples were collected after 1 h treatment with or without 1 mM NAA. E: Serial dilution assay illustrating the lethal phenotype of treatment with 1 mM NAA on cells expressing Mps1-AID-9xmyc in addition to OsTir1-9xmyc. The lethality can be rescued by expression of an exogenous Mps1 allele.

**Supporting Figure 4:**
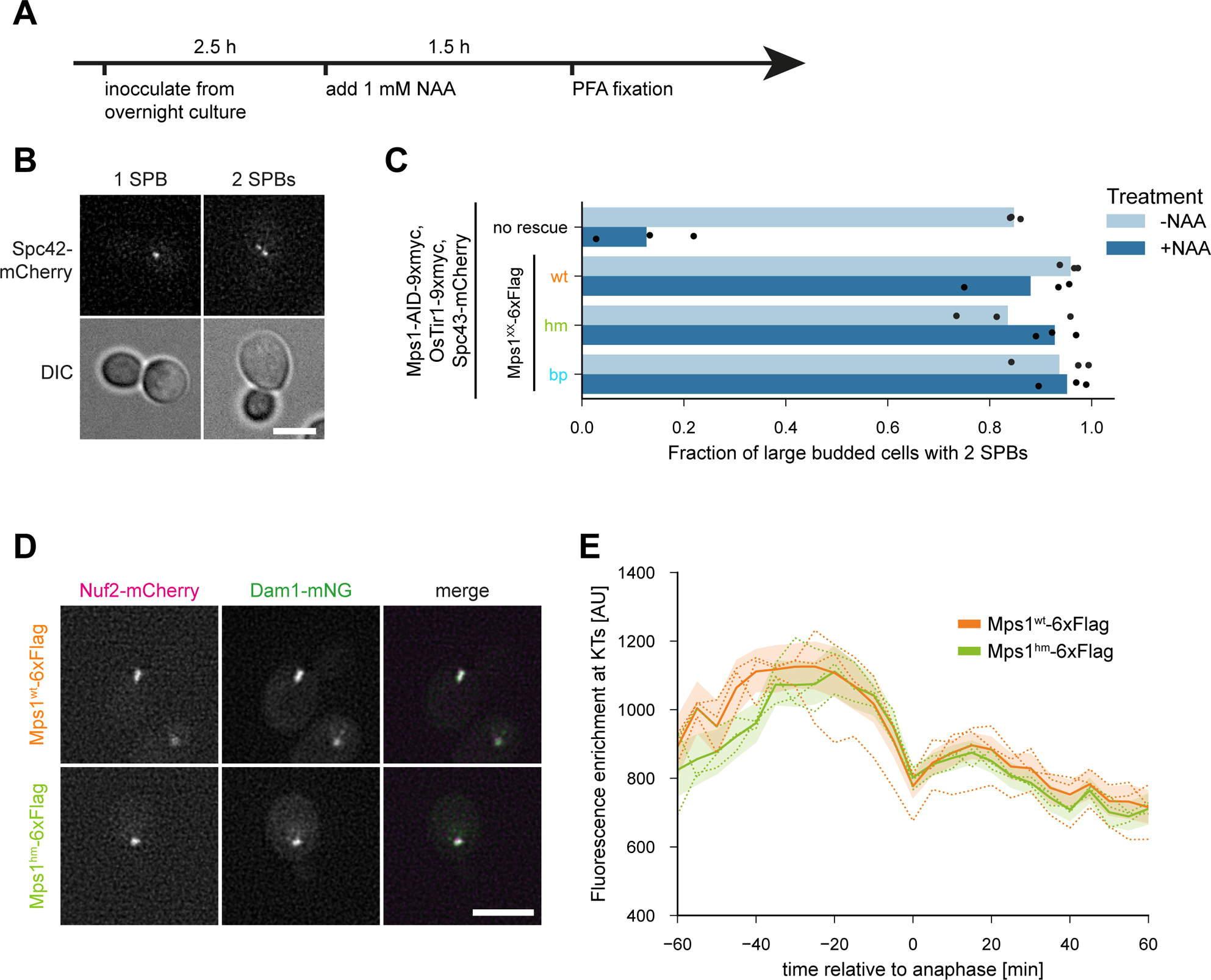
Further characterization of Mps1 interface mutants *in vivo*. A: Experimental scheme for the spindle pole body duplication assay. Logarithmically dividing cells were treated with 1 mM NAA to deplete the endogenous Mps1. Cells were harvested after 1.5 h, so large budded cells most likely entered the G1-phase after NAA addition. B: Example images for large budded cells displaying either one or two distinct signals for spindle pole bodies. Scale bar: 5 μm. C: Quantification of large budded cells showing two distinct signals for SPBs. Spindle pole body duplication is drastically impaired by depletion of Mps1 but can be rescued by expression of Mps1-6xFlag wildtype as well as the interface mutants. Bar graph: Mean fraction of three independent replicated. Dots: Fraction of the individual replicates. n>60 large budded cells per replicate. D: Representative micrographs of metaphase cells expressing Nuf2-mCherry and Dam1-mNeonGreen with Mps1^wt^-6xFlag or Mps1^hm^-6xFlag as their sole source of Mps1. No relevant difference is notable between cells expressing Mps1^wt^-6xFlag or Mps1^hm^-6xFlag. Scale bar: 5 μm. E: Quantification of the Dam1 fluorescence enrichment at kinetochores relative to the first anaphase frame. Mean and SEM of all replicates are indicated as continuous lines with borders, the means of three replicates are indicated as dotted lines.

## Notes

### Competing Interest Statement

The authors have declared no competing interest.

